# Extensively acquired antimicrobial resistant bacteria restructure the individual microbial community in post-antibiotic conditions

**DOI:** 10.1101/2024.08.07.606955

**Authors:** Jae Woo Baek, Songwon Lim, Nayeon Park, Byeongsop Song, Nikhil Kirtipal, Jens Nielsen, Adil Mardinoglu, Saeed Shoaie, Jae-il Kim, Jang Won Son, Ara Koh, Sunjae Lee

## Abstract

In recent years, the overuse of antibiotics has led to the emergence of antimicrobial resistant (AMR) bacteria. To evaluate the spread of AMR bacteria, the reservoir of AMR genes (resistome) has traditionally been identified from environmental samples, hospital environments, and human populations; however, the functional role of AMR bacteria in the human gut microbiome and their persistency within individuals has not been fully investigated. Here, we performed a strain-resolved in-depth analysis of the resistome changes by reconstructing a large number of metagenome-assembled genomes (MAGs) of antibiotics-treated individual’s gut microbiome. Interestingly, we identified two bacterial populations with different resistome profiles, extensively acquired antimicrobial resistant bacteria (EARB) and sporadically acquired antimicrobial resistant bacteria (SARB), and found that EARB showed broader drug resistance and a significant functional role in shaping individual microbiome composition after antibiotic treatment. Furthermore, longitudinal strain analysis revealed that EARB bacteria were inherently carried by individuals and can reemerge through strain switching in the human gut microbiome. Our data on the presence of AMR bacteria in the human gut microbiome provides a new avenue for controlling the spread of AMR bacteria in the human community.

## Introduction

Recently, the misuse and overuse of antibiotics in medicine and food production has led to the emergence of antibiotic-resistant bacteria^1, 2^. For example, the increased use of antibiotics during the COVID-19 pandemic has accelerated the development of multidrug-resistant (MDR) bacteria^3^, posing a new threat to modern society. Importantly, the human gut microbiome serves as a reservoir for antimicrobial resistance (AMR) genes^4–6^, spreading these genes to the community. Initially, monitoring the spread of AMR genes was conducted using environmental samples, such as urban sewage^7^, samples from hospital environments^8^, by studying normal human populations in different countries^9, 10^ and vertical transmission from mothers to infants^11^. Such surveillance approaches^12^ have been successful in the identification of AMR reservoirs, called resistomes, in the human community; however, a deep understanding of the taxonomical origins of AMR gene carriers and the impact of AMR bacteria on the gut commensal community is lacking. Some initial in-depth metagenomic studies have found that individual resistomes can persist for at least one year^9^, but the dynamic changes in AMR bacteria and their impact on the community have been poorly studied at species or strain resolution. Therefore, functional analysis of AMR bacteria in the individual gut microbiome will be a key to understand their functional niche in the human community and to control their unintended consequences in the human gut microbiomes.

Here, we performed a strain-resolved in-depth analysis of resistome changes using publicly available shotgun metagenomic samples of healthy adults after 4-day treatments with antibiotics cocktails^13^, focusing on resistome dynamics at the species/strain level through *de novo* assembly of the metagenome-assembled genome (MAG). We observed individual variations in AMR gene enrichment that tended to be concentrated within a few bacterial species. We categorized bacteria based on their AMR gene count into those that had acquired extreme resistance (EARB), those that had acquired sporadic resistance (SARB), and non-carriers, and compared their resistomes, taxonomy, and functional capabilities. EARB displayed broad drug resistance and a significant functional role in shaping individual microbiome compositions, suggesting that they may substantially alter the antibiotic-affected environment differently from commensal bacteria. Single nucleotide polymorphism (SNP) profile analysis revealed that multiple strains of EARB *E. coli* were inherently carried by individuals and reemerged through strain switching in the healthy human gut. We further confirmed the prevalence of EARB and its functional importance in additional cohorts with frequent antibiotic exposure, including those with recurrent urinary tract infections (RUTI) and liver cirrhosis, and preterm infants. Therefore, our findings provide insights into the distribution and enrichment of AMR genes within bacterial strains and highlight the functional significance of multi-resistant bacteria in the antibiotic-treated gut community.

## Results

### *De novo* assembly of shotgun metagenomics revealed the enrichment of multi-AMR genes within specific bacterial strains

To investigate the longitudinal changes in the anti-microbial resistance (AMR) gene repertoire of the gut microbiome after antibiotic treatment, known as the resistome, we obtained and processed publicly available fecal shotgun metagenomics data from 12 healthy adults. These data were collected before and after a 4-day treatment with antibiotic cocktails (meropenem, gentamicin, and vancomycin), as well as after the cessation of antibiotics at day 8, day 42, and day 180 (**Fig. 1**)^13^. First, we performed *de novo* genome reconstructions of 57 metagenomic samples using the metaWRAP pipeline^14^, which combines three representative metagenome binning tools: metaBAT2, CONCOCT, and maxBin2 (see **Methods**). A total of 7,858 initial bins of three binning tools were filtered into 2,585 high-quality (HQ) MAGs, which passed the quality criteria of completeness (> 70%) and contamination (< 5%) based on CheckM estimations^15^ (**Fig. 1b** and **Supplementary Fig. 1**a, b) and refined these into 1,358 MAGs, which were merged from the consensus bins of three binning tools (**Supplementary Table S1** and **Supplementary Data 1**).

**Fig. 1.**
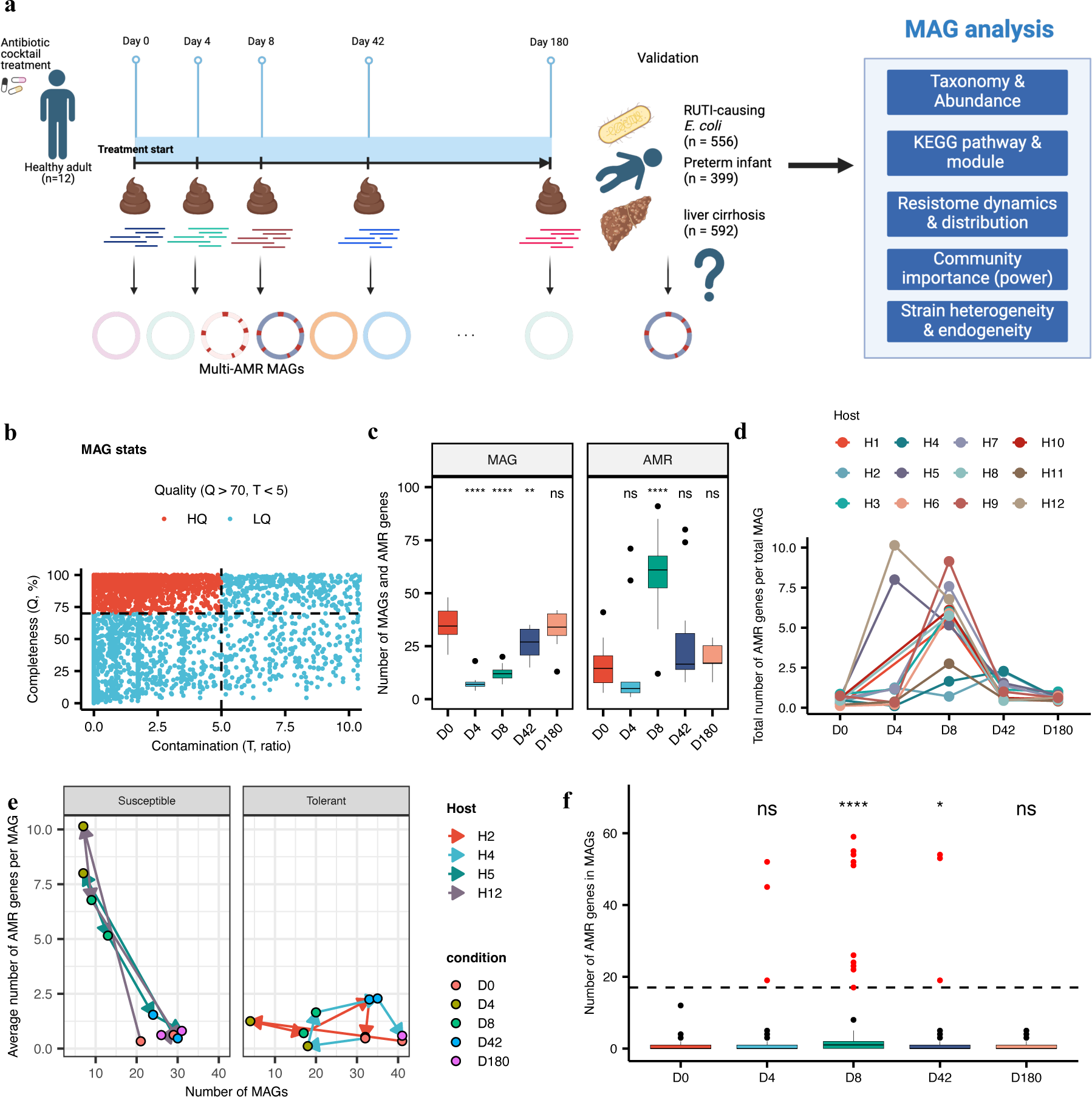
Metagenome-assembled genomes revealed individual resistome changes and discovered bacteria that had acquired extensive antimicrobial-resistance. **a,** Overview of the shotgun metagenome analysis conducted in this study. From 12 healthy adults, 56 metagenomics datasets were sampled at five different time points and three additional datasets (recurrent urinary tract infection [RUTI]-causing *Escherichia coli* genomes, preterm infant metagenomes, and liver cirrhosis metagenomes) were used for further validation. **b,** Scatter plot of metagenome-assembled genome (MAG) quality score of three different binning tools. Using CheckM algorithm, we selected high-quality MAGs (total number of high-quality (HQ) MAGs = 2,585 [blue], total number of low-quality (LQ) MAGs = 5,268 [red]) based on a completeness (Q) greater than 70% and contamination (T) less than 5% (block dotted lines). **c,** Boxplot of the number of *refined* MAGs based on the consensus of three binning tools (completeness > 70%, contamination < 5%; n = 1,358) and the number of antimicrobial- resistance (AMR) genes found in subjects at each sampling point. AMR genes were discovered by Resistance Gene Identifier (RGI) tool using the Comprehensive Antibiotic Resistance Database (statistical significance measured by Student’s t-test comparing each time point group with the day (D)0 group as reference group. \*\*\*\**p* ≤ 0.0001, \*\**p* ≤ 0.01). **d,** Individual dynamics of the number of AMRs per MAG. Each line indicates AMR changes in a given host. **e,** Line plots tracking the changes in the number of MAGs and average AMR gene burden per MAG of susceptible (Host 5 and 12) and tolerant subjects (Host 2 and 4). **f,** AMR gene distribution within each MAG (n = 1,358). MAGs containing more than 17 AMR genes (black dotted line) were regarded as bacteria that had acquired extreme resistance (EARB) (red dots, n = 20) (statistical significance measured by Student’s t-test comparing each time point group with the D0 group, \*\*\*\**p* ≤ 0.0001, \**p* ≤ 0.05).

We identified the AMR gene repertoire of the MAGs based on the homology of orthologous genes and SNPs using the Resistance Gene Identifier (RGI) tool of the Comprehensive Antibiotic Resistance Database (CARD)^16^. We traced the changes in the number of MAGs and AMR genes identified in each sample and summarized them by time point. The number of MAGs was highest at baseline (day 0), reduced significantly from day 4 to day 42, and then recovered on day 180. We also found a significant increase in the number of AMR genes by day 8, suggesting that 4 days after antibiotic treatment cessation, resistant bacteria persisted and thrived (**Fig. 1c**). Interestingly, individual examinations of changes in the AMR gene ratio revealed varied resistance responses of the gut microbiome to antibiotic treatments. For example, some individuals (host 5 and 12; susceptible group) showed drastic resistome changes from baseline, whereas other individuals (host 2 and 4; tolerant group) showed similar level of AMR gene burden, thereby implying different susceptibilities of individuals to antibiotic-induced dysbiosis (**Fig. 1d-e** and **Supplementary Fig. 1**c). In a previous study, administering antibiotics to healthy subjects not only altered their gut microbiomes to resemble those of intensive care unit patients, but also resulted in a protracted recovery in some subjects, as confirmed by the maximum displacement between baseline and final data points in principal component analysis.^17^ In line with these findings, our study revealed that individuals who experienced rapid enrichment of AMR genes following antibiotic treatment, i.e., susceptible group, demonstrated the longest Bray-Curtis distance from the baseline (day 0) to the last sample point (day 180) where showing recovered microbiome signature (distances among metagenomics samples by metagenomic-based operational taxonomic unit (mOTU) profiles) (**Supplementary Fig. 1**d).

Next, we checked the correlation between MAGs and the increased resistome by quantifying the prevalence of AMR genes within each MAG across time points (**Fig. 1f**). In our temporal analysis, we found that a specific set of 20 MAGs contained a notably high number of AMR gene, reaching a saturation point in the cumulative distribution of AMR counts, with each MAG containing at least 17 AMR genes (**Supplementary Fig. 2**a). Therefore, we defined bacteria with more than 17 AMR genes as extensively acquired antimicrobial resistance bacteria (EARB). The EARB showed comparable assembly quality compared to other MAGs (**Supplementary Fig. 2**b, c), indicating that the higher number of AMR genes in EARB was not an artifact of poor metagenome assembly. Of note, we found that individuals with different resistome changes, i.e., susceptible and resistant groups, showed different emerging timing of EARB species after antibiotics treatment (**Supplementary Fig. 1**c). Therefore, to better understand changes of individual microbiomes after antibiotics treatment, the dynamics of the abundance and functions of EARB species will be the key to understanding the effects of antibiotics toward gut microbiome community.

### Characteristics of EARB strains at the taxonomical, functional, and community level

We further explored the characteristics of EARB by comparing their taxonomy and abundance with those of other microbes in a given microbial community. We first annotated the taxonomy of all MAGs identified using Genome Database Taxonomy Toolkit (GTDB-TK)^18^ and constructed a phylogenetic tree for EARB (**Extended Fig. 1a**, see **Methods**). Of note, all EARB belonged to the Proteobacteria phylum (also known as Pseudomonadota), predominantly within the genera *Escherichia*, *Klebsiella*, *Enterobacter*, and *Cronobacter*, which are widely recognized as pathobionts associated with various chronic diseases and bloodstream infections^19^. We also checked the taxonomy of MAGs containing less than 17 AMR genes (termed SARB). The majority of SARB belonged to the phyla Bacteroidetes (also known as Bacteroidota) (62.8%) and Firmicutes (also known as Bacillota) (27.5%), whereas a few SARB belonged to Proteobacteria (3.8%). AMR non-carriers mostly belonged to Firmicutes (82.6%), implying different taxonomic preferences for the acquisition of antimicrobial resistance genes (**Supplementary Fig. 3**a, b).

Next, we estimated the relative abundance profiles of all MAGs at the phylum and family levels (**Extended Fig. 1b, c**). Interestingly, the abundance of *Enterobacteriaceae*, to which all EARB were assigned, and *Veillonellaceae*, known pathobionts associated with intestinal inflammation^20,21^, significantly increased when antibiotic-induced dysbiosis peaked (i.e. day 4 and 8). However, the abundance of the known commensal phyla, Bacteroidetes significantly decreased on day 4 and day 8 (Student’s *t*-test *p*-value = 0.0017 on day 8, compared with baseline). The pattern of changes in microbial composition was also confirmed using kraken2^22^ which showed resembled result (**Supplementary Fig. 3**c). We noted a shift in the microbial community composition within a short-term (day 0 to 42) following antibiotic treatment. This shift was characterized by a decreased relative abundance of EARB and an increased relative abundance of various minority species, suggesting a pattern of recovery from dysbiosis in the hosts (**Extended Fig. 1d** and **Supplementary Fig. 3**d).

### Differential resistomes between EARB and SARB strains

Next, we identified differences in the resistomes of EARB and SARB based on their annotated AMR gene information using the CARD^16^. Based on the resistome prevalence values (that is, the observed frequency of a given AMR drug class normalized according to the proportion of AMR-gene-carrying bacteria; see **Methods**), we compared the resistome composition differences at the drug class level between EARB and SARB strains (**Fig. 2a**). Of note, the resistome carried by EARB strains was more evenly distributed for many drug classes, whereas the resistomes carried by SARB were highly enriched for fluoroquinolone and tetracycline classes, which were not relevant to the antibiotics used in this study. Interestingly, a previous report showed that tetracycline-resistant genes are spread over many different geographical regions, which is the same tendency as that of the SARB resistome^23^. These distinct resistome between EARB and SARB was also confirmed in the pheatmap using relative frequency of drug class matrix (**Supplementary Fig. 4**a and b) and can be clustered again into EARB and SARB by unsupervised clustering (k-means) (**Supplementary Fig. 5**a). Among the three drug classes used in this study (carbapenem, aminoglycoside, and glycopeptide), the carbapenem resistance was found mostly in EARB (20 out of 23 MAGs of the given resistome were EARB) (**Supplementary Table S2**). Similarly, aminoglycoside resistomes were found mostly in EARB (20 of 40 MAGs of a given resistome were EARB). However, the glycopeptide resistome was found in SARB (44 MAGs), but not in EARB. The original study pointed out that gram-negative bacteria are naturally resistant to glycopeptide antibiotics, such as vancomycin, and EARB (all gram-negative) might have natural resistance to such glycopeptide antibiotics, thereby not necessitating a glycopeptide resistome^24^.

**Fig. 2.**
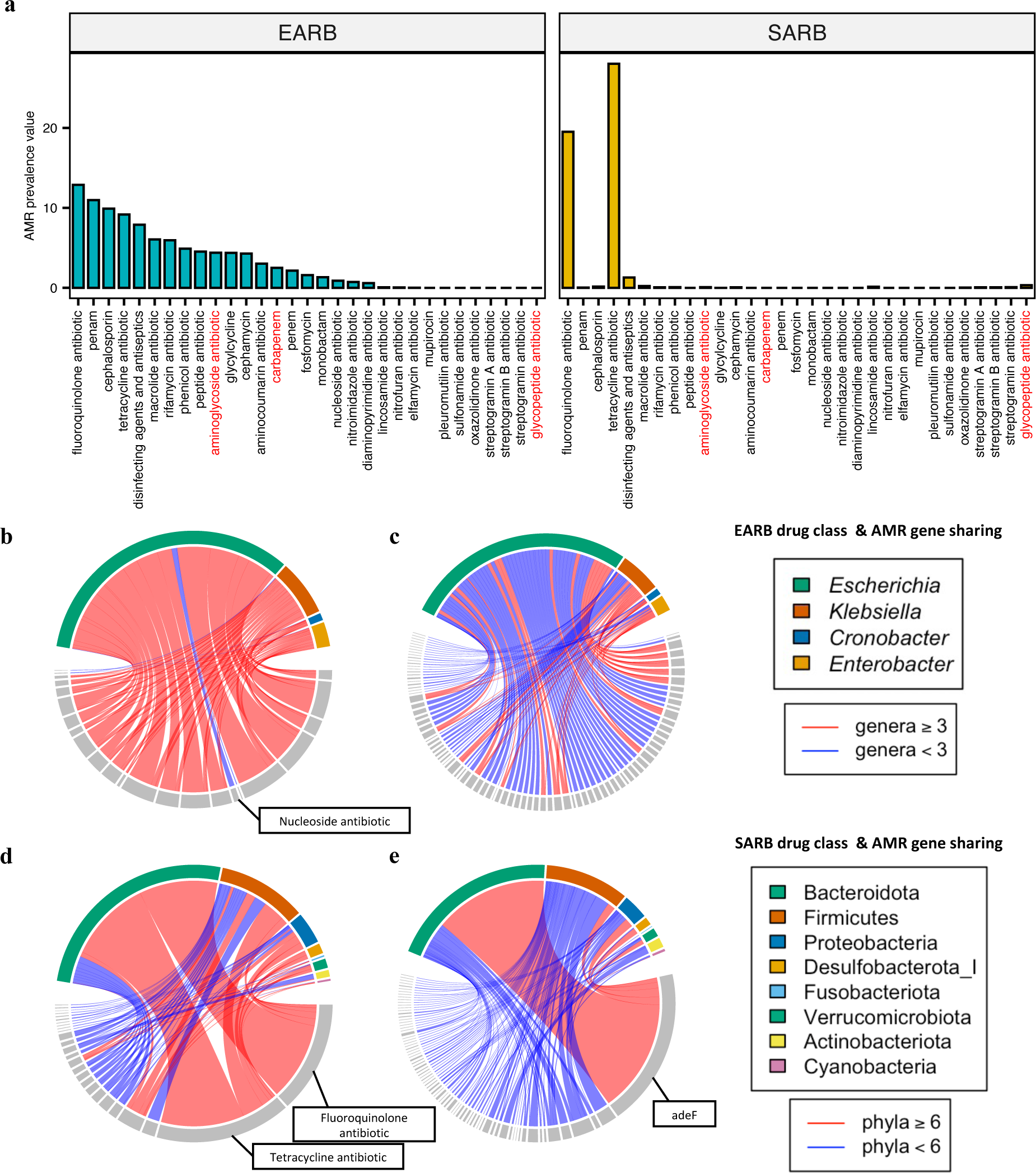
Differential resistomes between EARB and SARB strains. **a,** drug classes resistant to EARB and SARB strains identified by AMR prevalence scores. We calculated AMR prevalence scores based on the observed frequency of given drug classes in EARB or SARB strains, while normalizing them according to their carrier ratios. We show the AMR prevalencescores from EARB (left) and SARB (right) strains based on the descending orders of EARB prevalence scores (the class of antibiotics used for treatment in this study are colored red). **b- c,** Circos plots of (**b**) shared drug classes (n = 23) and (**c**) shared AMR genes (n = 803) (upper bounds) of given genera of EARB (n = 4) (gray, lower bounds). Red links indicate drug classes or AMR genes shared by more than three genera, whereas blue links indicate those shared by less than three genera. **d-e,** Circos plots of (**d**) shared drug classes (n = 26) and (**e**) shared AMR genes (n = 832) of given phyla of SARB (n = 8). Red links indicate drug classes or AMR genes shared by more than six phyla, whereas blue links indicate those shared by less than six phyla.

We further investigated the resistome profiles of EARB and SARB based on the shared AMR drug classes (**Fig. 2b, d**). Of note, all EARB shared resistance to similar antimicrobial drug classes, whereas SARB did not, except for fluoroquinolone and tetracycline. Therefore, the extensive antibiotic resistance capacity of EARB provides a fitness advantage when treated with antibiotics, whereas SARB show different survival outcomes according to the type of antibiotic used. However, at the gene level, neither the EARB nor the SARB strains shared AMR genes with other phylogenetic groups (genera or phyla), implying that similar drug class resistomes were conferred by unique or narrowly distributed AMR genes (**Fig. 2c, e**). This result also suggests that horizontal gene transfer (HGT) of AMR genes between bacteria of different taxa is uncommon. Rather, a specific taxonomic group harboring an extensive resistome and AMR gene spread through HGT might only occur in specific genes.

We also traced the changes in resistome profiles at different time points after extracting resistome signatures using non-negative matrix factorization (NMF) (**Supplementary Fig. 6**a, see **Methods**). By applying an unsupervised NMF decomposition to the resistome profiles, we identified two distinct signatures in the resistome, *sig1* and *sig2*. As expected, these two signatures resembled the EARB and SARB resistome prevalence scores, thereby confirming our findings using independent methods. Of note, EARB-like resistome signature (*sig1*) was significantly increased on day 4 and 8, whereas SARB-like resistome signature (*sig2*) was decreased on day 4 and 8 and then recovered at day 42 and 180. Such resistome signatures imply that EARB drives overall resistome changes after the antibiotics treatment, as observed in the data presented in **Fig. 1**.

### EARB strains showed functional dominance in the post-antibiotic gut microbial community

In our previous findings (**Extended Fig. 1** and **Supplementary Fig. 3**), we identified EARB strains co-enriched with specific bacterial groups, suggesting that EARB play a significant role in shaping a given community when treated with antibiotics. Therefore, to further investigate the impact of EARB within the community, we devised a new scoring method that estimates functional dominance, termed as the *community power score* (**Fig. 3a**). In short, the community power of a given microbe was calculated by summing the proportion values of its genes associated with various functions (i.e., Kyoto Encyclopedia of Genes and Genomes [KEGG] orthologs), which reflected the microbe’s functional uniqueness and importance within the community (see **Methods**). Remarkably, we found that, in all metagenomic samples, EARB showed the highest community power compared to the other microbes, indicating its significant functional importance on the community. However, SARB and AMR non-carriers showed significantly lower community power scores than EARB strains (Student’s *t*-test *p*-values < 1 × 10^-4^) (**Fig. 3b**, right).

**Fig. 3.**
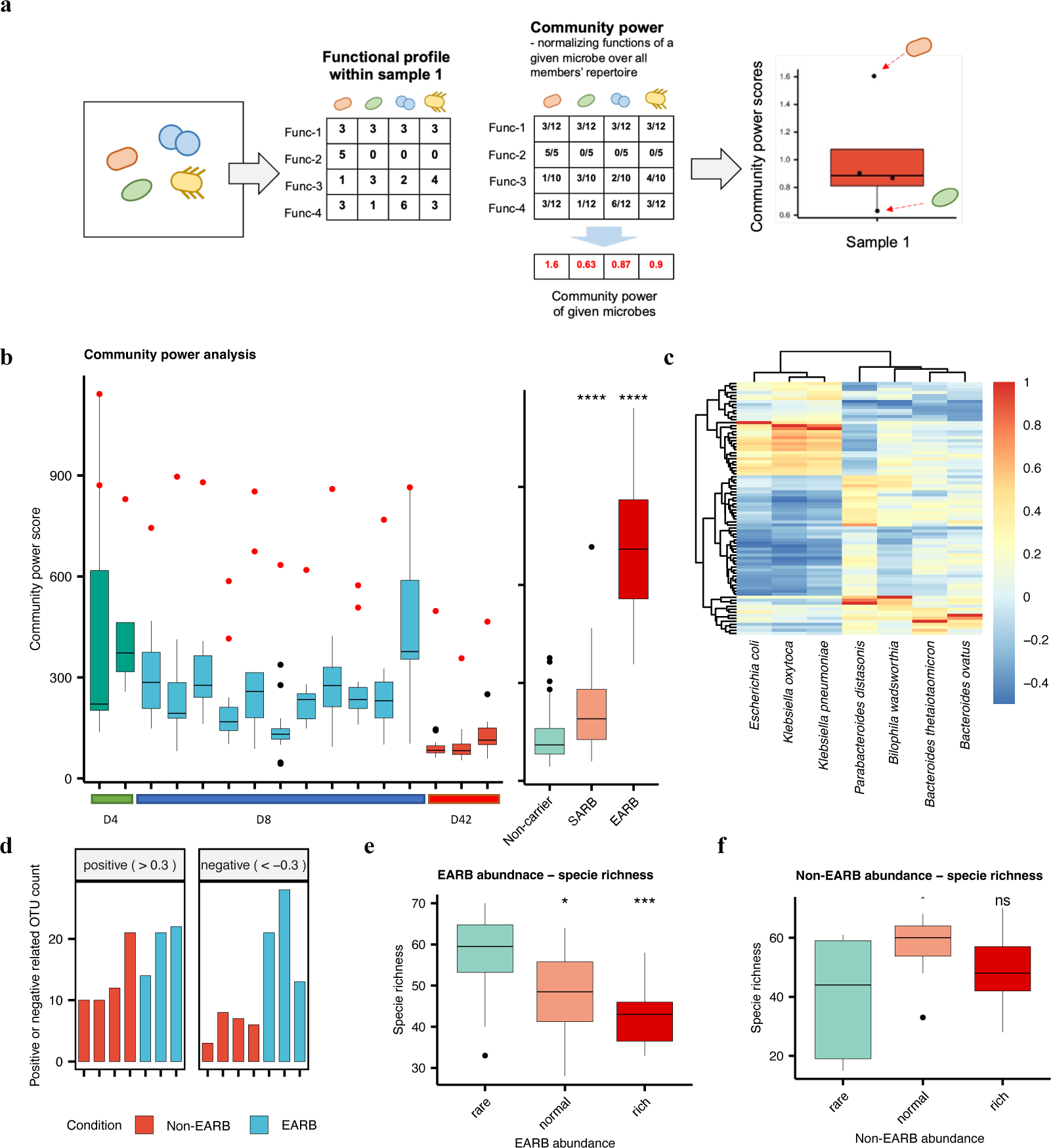
EARB strains harbored a greater functional repertoire capable of shaping the community. **a,** The outline of community power analysis. We calculated the functional repertoire of given microbial strains, herein termed community power scores, based on the number of genes harboring specific Kyoto Encyclopedia of Genes and Genomes orthology (KO) terms. In short, we counted the total functional repertoire of given microbial strains based on their genes harboring specific KO terms, after normalizing them according to the total number of KO terms in all the microbial strains in given metagenomic samples. Therefore, all community power scores for microbes in a given metagenomic sample could be calculated and compared with each other. The microbes of the highest community power score had the highest functional repertoire in a given metagenomic sample. **b**, Community power scores of metagenomic samples that carried EARB strains (n = 16). We compared the community power scores among EARB, SARB, and non-carriers (right panel) and found that EARB outscored the others (Student’s t-test *p*-values < 0.0001 [****]). **c**, Co-abundance (i.e., Spearman’s correlation coefficients) of three EARB strains (*Escherichia coli*, *Klebsiella oxytoca*, *Klebsiella pneumoniae*) and four non-EARB strains with the highest community power score in one of the metagenomic samples (*Bacteroides thetaiotaomicron*, *Bacteroides ovatus*, *Bilophila wadsworthia*, *Parabacteroides distasonis*). Microbial strains significantly correlated with at least one of the seven selected strains (absolute correlation coefficients > 0.3) are shown and unknown metagenomic-based operational taxonomic unit(mOTU) species were excluded. Additionally, mOTUs that were not present in more than five metagenomic sample were also excluded (rows in the heatmap, n = 84). **d,** The number of significantly correlated strains (absolute Spearman’s correlation coefficients > 0.3) with EARB and non-EARB strains. **e,** Negatively correlated between EARB abundance and species richness, based on the sum of log2-transformed relative abundances (rare < -10, -10 ≥ normal < -5, rich ≥ -5) and species richness. **f,** Less-correlated patterns between sample groups of different non-EARB abundances, based on the sum of log2-transformed relative abundances (rare < -10, -10 ≥ normal < -5, rich ≥ -5) and species richness (relative abundance > 0). Differences between samples groups were tested using Wilcoxon rank-sum tests.

To further identify which community members were influenced by EARB strains, we checked for microbes that were co-abundant with EARB (*Escherichia coli*, *Klebsiella oxytoca*, and *Klebsiella pneumoniae*) and other non-EARB (i.e., SARB and non-carriers) with the highest community power scores (*Bacteroides thetaiotaomicron*, *Bacteroides ovatus*, *Bilophila wadsworthia*, and *Parabacteroides distasonis*) (|Spearman’s correlation coefficients| > 0.3) based on mOTU-based species abundance profiles (see **Methods**). Interestingly, co-abundant EARB were mostly pathobionts (**Supplementary Table S3**), including *Clostridium* spp., *Fusobacterium* spp., and *Veillonella* spp., which promote intestinal inflammation^20^. In contrast, members that were co-abundant with non-EARB were mostly commensal bacteria, including *Bacteroides* spp., *Roseburia* spp., and *Alistipes* spp. When we checked the negative bacterial interactions between EARB and non-EARB, we found that EARB showed a much higher number of negatively correlated microbes than non-EARB, implicating EARB outcompeting in given microbial community (**Fig. 3d**). We substantiated our findings using correlations between EARB abundance and species richness and identified negative trends between these two parameters (**Fig. 3e**). However, the abundance of non-EARB with high community power correlated with increased species richness in the middle-range abundance level (**Fig. 3f**). Therefore, we identified two distinct patterns of functionally impactful bacteria – 1) microbes with the high community power that might influence a small set of bacteria and 2) microbes with high community power that are co-abundant with a large number of bacteria, behaving like commensal species.

Next, to better characterize the functional impact of EARB on other interacting microbes, we performed functional enrichment analysis of EARB compared to non-EARB based on the KEGG pathways and modules^25^ (Wilcoxon rank sum test *p*-values < 0.05, log2 fold change > 0.5, pathway coverage > 0.3 or module coverage > 0.8; **Fig. 4a and b**). Interestingly, we found that EARB were significantly enriched in amino acid metabolism; sugar metabolism; and reductive transformations of xenobiotics, including reductions in aromatic compounds, such as processes supplemented with enriched co-factor biosynthesis (e.g., glutathione, vitamin B6, and ascorbate). Notably, xenobiotic degradation is known to support anaerobic respiration through nitrogen and sulfur reduction, promote biomass generation of the given microbiota, and provide a fitness advantage^26^, and is also likely to provide the transformation capacity for drug compounds, including antibiotics. In addition, we found multiple biosynthesis pathways for antibiotic-like compounds in EARB, which may include bacteriocin, a narrow-spectrum antimicrobial substance^27–29^. In summary, we found that the EARB strains were equipped with biosynthetic pathways, xenobiotic-reducing metabolism, and anaerobic respiration capacity.

**Fig. 4.**
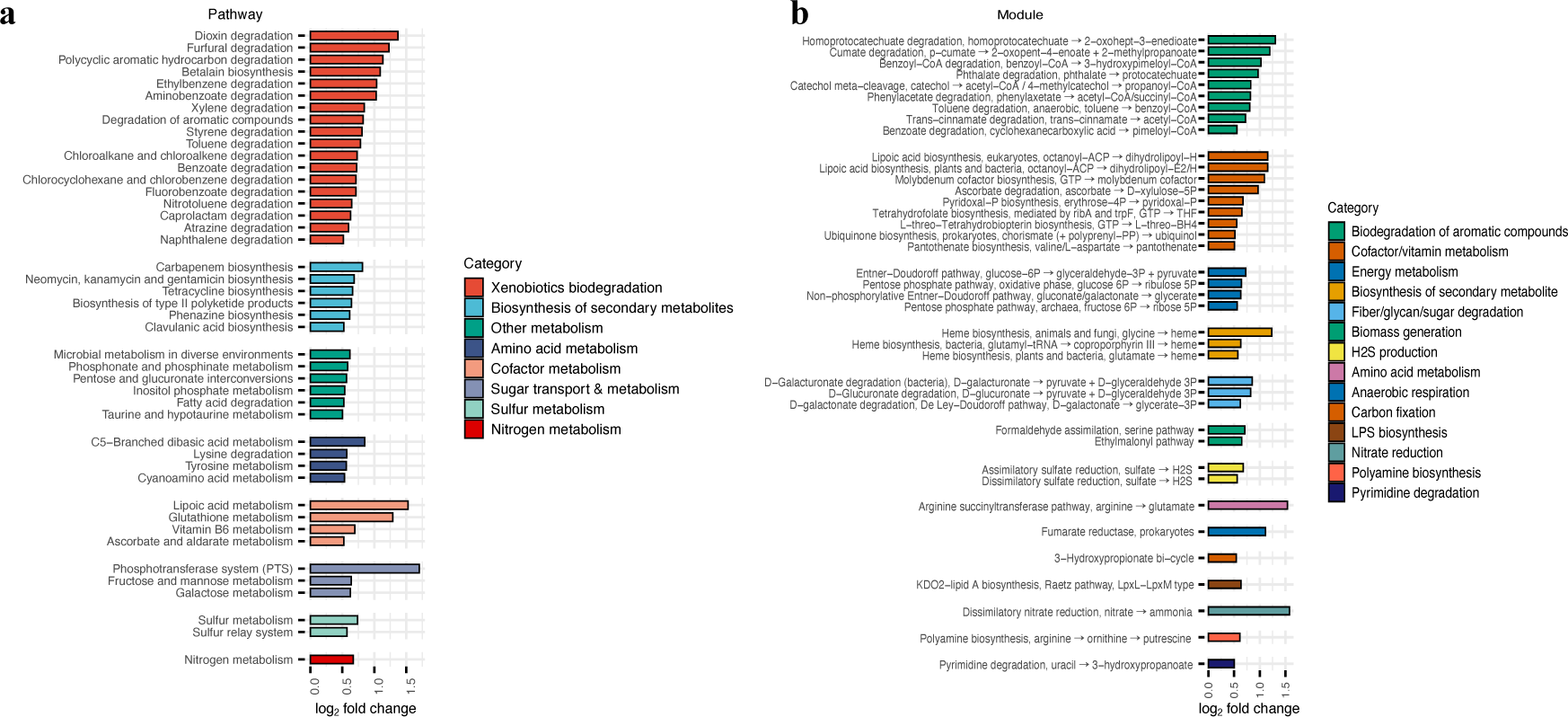
The enriched metabolic pathways of EARB strains in a given community. **a-b,** (**a**) KEGG metabolic pathways or (**b**) KEGG modules significantly enriched among EARB strains compared to other microbes in given metagenomic samples (total number of EARB = 20 and total number of non-EARB compared = 217, Wilcoxon rank-sum test *p*-values < 0.05, log-2 fold change > 0.5, and pathway coverage > 0.3 or module coverage > 0.8).

### Multiple EARB strains were carried endogenously and alternated in individual hosts

Next, we checked EARB strain changes after antibiotic treatment to determine whether they were carried endogenously from the same hosts or re-colonized spontaneously from outside the host. To this end, we first selected EARB strains from different hosts (n = 20) and checked their genomic similarities by the average nucleotide identity (ANI) scores^30^ (see **Methods**). The ANI scores showed distinct patterns between *E. coli* and other EARB of different genera (**Supplementary Fig. 7**a). When we focused on the ANI scores between the EARB *E. coli* strains (n = 12), we observed considerable strain diversity, ranging from 96.56 to 99.99% (**Fig. 5a**). Of note, EARB *E. coli* strains from the same host (i.e., host 12) on day 4 and 8 showed a substantially high ANI score of 99.99% (**Fig. 5b**). Given that identical strains were defined based on an ANI score of 99.8%^31^, the two EARB *E. coli* strains from the host 12 could be regarded as identical, implying same strains endogenously carried between day 4 and day 8. In contrast, EARB *E. coli* strains from host 5 on day 4 and 8 showed distinct ANI scores (99.40% and 98.78%) (**Fig. 5b and Supplementary Fig. 7**b). When comparing with other EARB *E. coli* strains, host 12 *E. coli* strains showed identical ANI scores whereas host 5 *E. coli* strains showed different ANI score patterns. Therefore, we found substantial strain heterogeneity of EARB strains of *E. coli*, even between those originating from the same hosts.

**Fig. 5.**
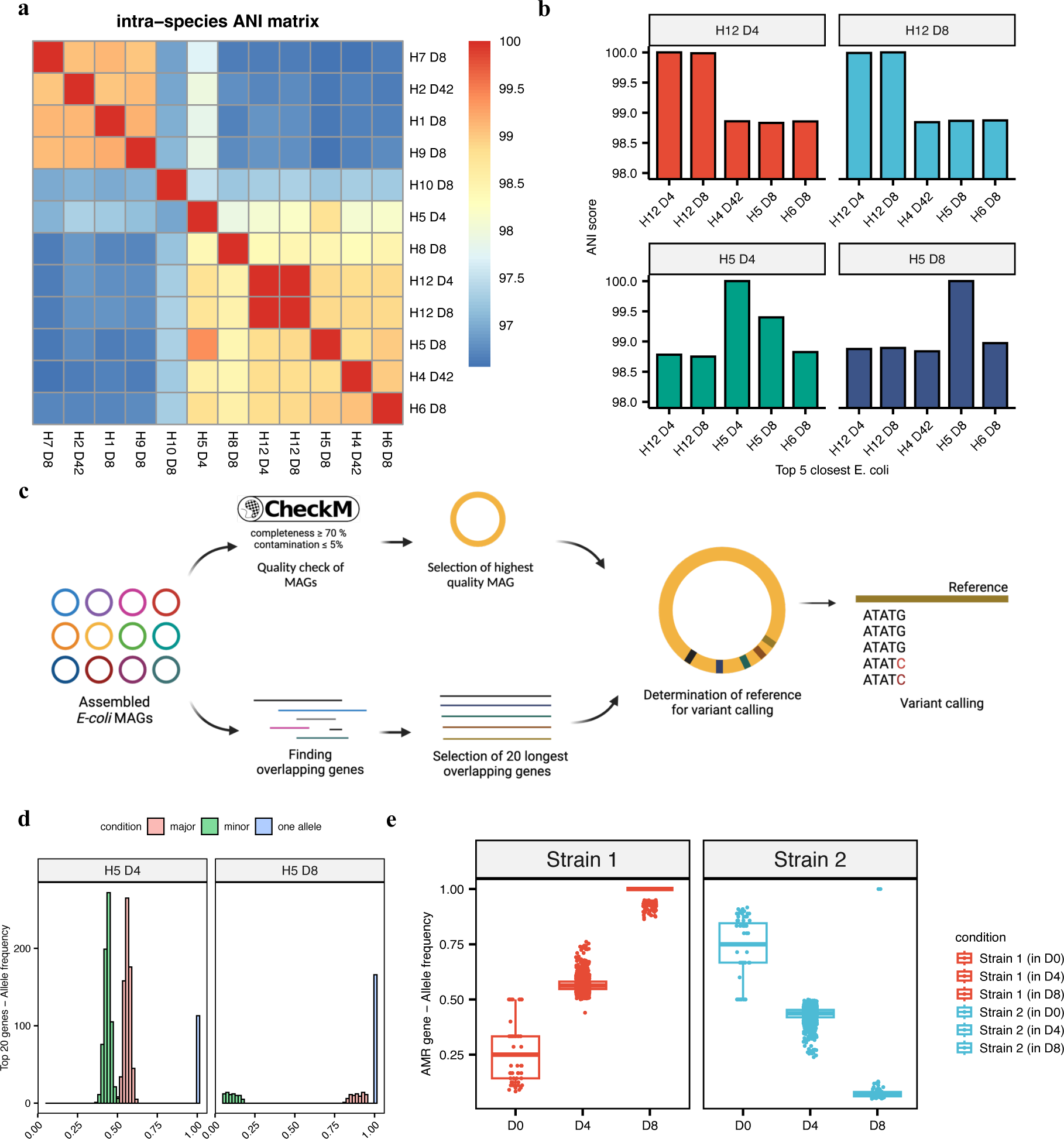
EARB strains were carried endogenously by some hosts and major strains alternated after antibiotic treatment. **a**, Intra-species average nucleotide identity (ANI) scores of *E. coli* strains belonging to EARB (n = 12). **b**, Top 5 closest EARB *E. coli* strain ANI scores with two specific strains identified from the same host (H5, H12). **c**, The workflow for the identification of single nucleotide polymorphisms (SNPs) from a given metagenome sample. First, we selected 20 homologous genes with the longest lengths from all EARB *E. coli* MAGs as the on-target genes for SNP analysis. Next, we used the 20 gene set from the best-quality MAG for building reference and performed variant calling for metagenomics samples which contain EARB *E. coli* strains. **d**, Allele frequency distributions of SNPs identified in metagenomic samples of host 5 at day 4 (left, n = 805) and day 8 (right, n = 240). We identified two different patterns of major and minor alleles, and also one allele existing in both strains. **e**, SNP analysis was conducted using *E. coli* AMR genes of host 5 at day 8 as a reference set. We found that two *E. coli* strains shared same variant positions and different SNP profiles (nucleotide) under different conditions (median of allele frequencies for strain 1 at day 0, 4, and 8 = 0.250, 0.563, 1.000, respectively and those for strain 2 at day 0, 4, and 8 = 0.750, 0.437, 0.0694, respectively).

To further identify strain changes in the longitudinal samples, we performed SNP calling of homologous genes present in host 5 EARB *E. coli* strains (**Fig. 5c** and **Supplementary Table S4**, see **Methods**). Briefly, we identified 20 homologous gene, possessed by all *E. coli* strains, then built reference using homologous gene set of best quality *E. coli* strains (H6 D8). Next, we called the variant positions (SNPs) by mapping the host 5 metagenome reads and checked the shared variant positions and calculated allele frequencies of all variant positions. As a result, we found 804 and 240 SNPs of homologous gene reference from day 4 and 8, respectively (**Fig. 5d**). Analysis of the allele frequency of the identified variants revealed two separate clusters on day 4, with the major cluster being close to 55% and the minor cluster being close to 45%. However, on day 8, one allele converged to almost 100% (median: 98%). This suggests that one of the two strains present on day 4 became dominant on day 8.

Since, we identified the existence of two distinct strains in host 5, we tried to track down the changes of strains using EARB *E. coli* AMR gene set of host 5 on day 8 (number of AMR genes = 54) as a reference due to the high dominance of single strain. Our analysis identified 1,841 variants on day 4 across 49 AMR genes out of 54 AMR reference gene sets. Considering that a single strain was dominant on day 8, while two strains were present on day 4, we hypothesized that the variants identified on day 4 were due to the presence of a minor strain. Consequently, the five genes without variants may either be due to perfect matching in either strain or may only be found in the major strain. Examination of the read depth of the AMR genes revealed that one of the five genes presented a depth similar to that of the other 49 genes carrying variants, suggesting that it was shared between the two strains and perfectly matched. The remaining four genes displayed significantly lower depths, indicating that they were exclusive to the major strain (**Supplementary Fig. 7**c). Allele frequency analysis of the 1,841 variant positions on day 4 revealed a distribution similar to that of the homologous genes, with approximately 55% major strains and 45% minor strains (**Fig. 5e and Supplementary Table S5**). Comparing the major and minor nucleotides of these variant positions longitudinally by tracking same variant position with same SNP nucleotide between day 4 and day 8, we found that the major strain on day 4 became dominant on day 8. To determine whether these AMR genes were also present at baseline, we also tracked SNP profile on day 0 (**Fig. 5e**). Although only a single allele was confirmed at many variant positions owing to the shallow sequencing depth (single allele position = 333), the major and minor strains were still identifiable at certain positions (two-allele position = 76). Remarkably, the strain that was predominant on day 8 was observed as a minor strain on day 0, suggesting that the emergence of this EARB may be attributed to strain alterations. As the endogenous nature of these *E. coli* strains has been established in a previous study^32^, our results confirmed that the occurrence of EARB in an antibiotic-exposed environment is caused by opportunistic pathogenic bacteria that already inhabit the host.

### EARB strains identified in diverse antibiotic-induced dysbiotic conditions

We further investigated other populations with frequent antibiotic exposure to determine if their microbiota contained EARB (**Fig. 6**). For example, we selected shotgun metagenomic cohorts and the genomes of bacterial isolates showing antibiotic resistance, such as those causing RUTI^32^, and performed *de novo* assembly to identify EARB strains harboring more than 17 AMR genes (see **Methods**). First, we found that all *E. coli* isolates from RUTI were EARB strains (mean number of AMR genes it carries = 61; **Supplementary Table S6**). Interestingly, based on SNPs that defined major and minor strains in a previous analysis (**Fig. 5**), we found that most *E. coli* isolates (65%) resembled the major strain’s SNP profile (i.e. having more than half of same nucleotides of 1,841 SNPs that define major strains; **Supplementary Fig. 8**), whose abundance was increased by antibiotics. This implies that, even within RUTI-EARB *E. coli*, the SNP patterns of the major strain were more prevalent in antibiotic-exposed environments. Next, we checked other MAGs assembled from antibiotic-exposed cohorts (i.e., preterm infant^33^ [n = 399] and liver cirrhosis cohorts^34^ [n = 592]) and found EARB strains among the selected cohorts (303 of 399 infant samples and 60 of 592 liver cirrhosis samples) (**Fig. 6**). More interestingly, when we checked their resistome profiles (**Fig. 6a**), we found that they showed highly similar resistome profile to the EARB strains we found in a previous analysis (**Fig. 2a**). Based on further functional investigations, such as community power scores (**Fig. 6b**), we found that EARB had the greatest functional impact on a given community likewise. Of note, preterm infants showed a higher prevalence of EARB strains, whereas patients with liver cirrhosis had a higher average number of assembled MAGs than preterm infants (**Fig. 6c**, **Supplementary Table S7, 8**), implying that preterm infants are more susceptible to EARB colonization, as they have a simple microbial community with less colonization resistance^35^. Additionally, metagenomic analysis of samples from patients with liver cirrhosis from two different samples, stool (n = 264) and saliva (n = 328), showed that the prevalence of EARB strains was higher in stool samples (n^stool^ = 57 and n^saliva^ = 7). These results showed that EARB prevalently exist within population who treated antibiotic multiple times and show same characteristics with our finding. Therefore, we confirmed the existence of EARB in diverse populations based on the newly discovered MAGs from different data sources (i.e., isolate genomes, single-end and paired-end sequencing data).

**Fig. 6.**
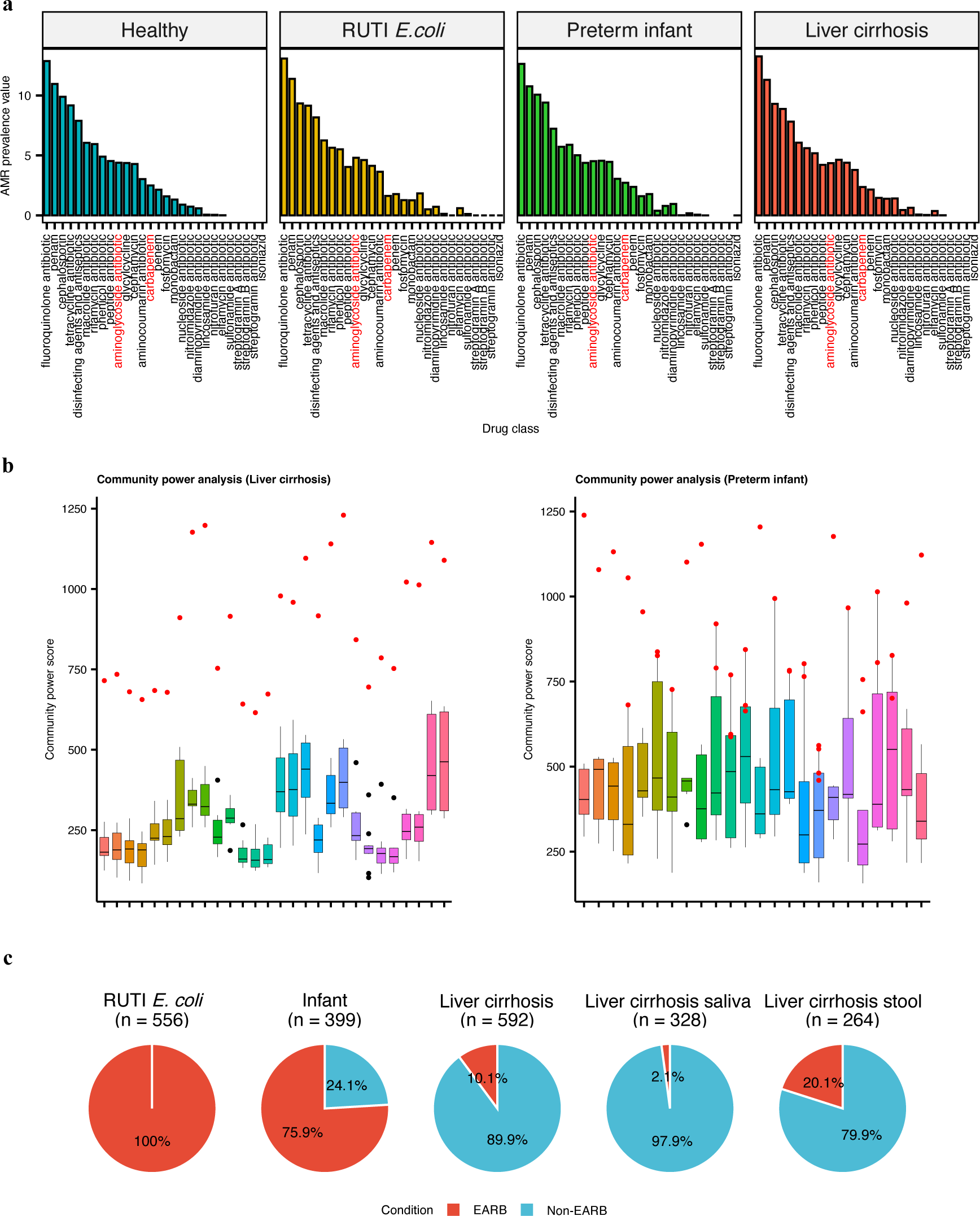
EARB strains found in different antibiotic-exposed cohorts with the greatest functional impact on the community. **a**, AMR prevalence comparison between data from healthy adults and data from three different cohorts (recurrent urinary tract infection [RUTI]-causing *E. coli*, preterm infants, and patients with liver cirrhosis). **b**, Community power analysis applied to preterm infants, and patients with liver cirrhosis). **b**, Community power analysis applied to liver cirrhosis (left, n = 592) and preterm infant (right, n = 399) study data. Community poweranalysis was conducted on samples harboring five MAGs and at least one EARB strain (red dot indicates the EARB strain). **c**, Proportion of samples containing EARB strains in three different datasets (*E. coli* isolate, preterm infant, and liver cirrhosis [saliva and stool]).

## Discussion

The need for research on antibiotic-resistant bacteria at the species and subspecies levels has increased. In this study, we performed *de novo* assembly of MAGs to investigate AMR gene dynamics at the species and strain levels within microbiomes altered by antibiotics. First, using the MAG-based approach, we found different responses of individual microbiomes to antibiotics, which were classified into susceptible and tolerant groups. Second, based on AMR gene repertoire analysis, we identified two distinct groups of antibiotic-resistant bacterial strains in individual microbiomes: EARB and SARB. Of note, pathway enrichment analysis showed that EARB harbored specific functions that enhanced their survival and competitive advantage, such as the degradation of xenobiotic substances, including aromatic antibiotics, and the promotion of biosynthetic pathways. Third, we found that EARB harbored a wide range of AMR genes for different drug classes compared to SARB, which were mostly enriched for only tetracycline and fluoroquinolone antibiotic resistance. Fourth, we found that EARB played a significant role in altering post-antibiotic microbial community dynamics based on the community power score we devised. Finally, we identified multiple resistant lineages of EARB strains and their intra-species competition within the same host during antibiotic treatment, highlighting the importance of species-specific strategies to manage antibiotic resistance.

Notably, functional profiles of MAGs indicated enhanced xenobiotic degradation pathways within the EARB, suggesting their potential role in the persistence of antibiotic-susceptible bacteria via degradation enzymes, such as beta-lactamases. In addition, we identified enrichment of secondary metabolites or antibiotic-like compound biosynthesis pathways among the EARB strains. This suggests a competitive advantage for the EARB strains, facilitating their survival in the intestines of healthy individuals^36^. These findings elucidate the complex interplay between EARB and the gut microbiota, potentially driving the maintenance of antibiotic resistance. Therefore, further functional analysis is essential to fully understand how EARB promotes the co-enrichment of antibiotic-susceptible bacteria and how they promote their persistence in the intestines of healthy individuals.

In addition, we performed strain-level analysis of the EARB of individual microbiomes, particularly *E. coli*, and identified remarkable genomic diversity within strains isolated from the same host over time, that is, the existence of multiple strains or lineages. Longitudinal data analyses allowed us to track the development of these EARB *E. coli* strains, uncovering two key patterns: the presence of native strains within the host gut and the occurrences of a “strain sweep”, where certain strains outcompete others, potentially due to superior resistance capabilities or other fitness advantages^37^. This advantage can also be explained by the observed weighted SNP patterns in RUTI-causing *E. coli*, suggesting that these differences may influence the adaptability of bacteria to antibiotic environments. These findings indicate that the gut is a reservoir for diverse antibiotic-resistant strains and demonstrate the dynamic nature of microbial evolution influenced by selective pressures. Such observations have significant implications for therapeutic strategies, highlighting the need for in-depth research to elucidate the determinants of strain variability and enhance approaches to effectively manage antibiotic resistance.

In summary, our study offers a comprehensive investigation of the dynamics of antibiotic-resistant bacteria through MAG analysis. Our MAG-based approach provided a detailed understanding of the entire resistome at the individual species level. Here, we found that EARB became a dominant force in microbial communities following antibiotic treatment, and that their genetic diversity and evolutionary pathways present notable challenges in managing antibiotic resistance. These findings not only deepen our understanding of antibiotic resistance dynamics but also pave the way for more targeted strategies to combat the spread of resistant bacteria.

## Methods

### De novo Metagenomic Assembly

We obtained high-quality non-human reads from original paper^13^. Human contaminations were removed from metagenomic samples again using GRC38 and bowtie2 (version 2.3.5.1)^38^. Host removed reads were used for MAG construction using metaWRAP (version 1.3.2) pipeline^14^. First, FASTQ files were assembled by MEGAHIT (v1.1.3)^39^. Assembled files were binned by three different tools (MetaBat2, MaxBin2, CONCOCT). Initial bins were refined to get final MAG. CheckM (version 1.0.12)^15^ was used for quality control for MAG and MAGs with >70% completeness and <5% contamination were further considered as high-quality.

### Antimicrobial resistance gene analysis

Antimicrobial resistance (AMR) genes on assembled genome were searched by Resistance Gene Identifier software (RGI; version 5.2.1) with CARD database (3.2.2)^40^ as a reference (-- input_type contig --alignment_tool DIAMOND --split_prodigal_jobs –clean). MAGs that contain more than 17 AMR genes were defined as Extremely Acquired Resistant Bacteria (EARB) in this study and MAG containing AMR genes less than 17 were considered as Sporadically Acquired Resistant Bacteria (SARB); MAGs having no AMR genes were considered as non-carrier.

To analyze AMR dynamics, we counted the number of MAGs and AMR genes at different sampling points. In addition, we tracked the average number of AMR genes per MAG (by dividing the total number of AMR gene by the number of MAG) to identify individual variations in response to antibiotic treatment. We categorized individuals into ’Susceptible’ and ’Tolerant’ groups by considering the timing of peak of AMR gene increases and also degree of the increase. To confirm microbial composition displacement among the tolerant group, we used mOTU relative abundance, from the original paper^13^, and R package *vegan* (version 2.6- 4) to calculate Bray-Curtis dissimilarity between day-0 (baseline) and day-180 of all subjects. We obtained multi-dimensional scaling (MDS) coordinates and calculated the displacement. Drug classes and mechanisms of AMR genes in each MAG were obtained from RGI analysis result. To analyze the distribution of various drug classes, number of occurrences of drug classes were counted in EARB and SARB. To consider the prevalence of drug classes along EARB and SARB, the frequency of each drug class was calculated and number of MAGs that carried each drug class was counted. To get a prevalence score, proportion of MAG carrying a given drug class (carrier ratio) were multiplied with frequency of the drug class.

To analyze sharing of drug classes and AMR genes among EARBs and SARBs, we counted the frequency of drug class or AMR gene at certain taxonomic level (genus level for EARB and phylum level for SARB). We considered the cases of EARB as sharing if EARBs included in 4 genus or if a drug class or AMR gene was found in more than 3 genera. On the other hand, we considered the cases of SARB as sharing if SARB included 8 phylum and if a drug class or AMR gene was found in more than 6 phyla.

To explore potential correlations between specific drug classes or AMR genes and taxonomies, we calculated the frequency of each drug class within every phylum or species. This was done by dividing the number of times an AMR gene associated with a drug class appeared within a given taxonomy by the total occurrences of that taxonomy level. We performed this calculation separately for EARB and SARB. Following the frequency determination, we added one to the mean frequency value to adjust for any instances of zero occurrence, then applied a logarithmic transformation to normalize the data distribution. We visualized the results using a pheatmap to provide an intuitive understanding of the frequency and distribution patterns. We also checked the AMR signature from MAGs based on non-negative matrix factorization (NMF) technique using R package *NMF*. By setting factorization rank as two, we extracted basis and coefficient vectors from given NMF factors obtained.

### Taxonomy annotation of MAGs

The taxonomy of MAGs, assembled in this study, was assigned using GTDB-TK classify_wf (version 2.1.0) with release 207_v2 database^18^. Phylogenetic tree for EARB constructed from 12 healthy adults was constructed also using GTDB-TK de_novo_wf. Since all EARBs were assigned within *Enterobacteriaceae* family, following parameter was used (--bacteria -- taxa_filter o Enterobacterales --outgroup_taxon f *Enterobacteriaceae*). The result was used for plotting phylogenetic tree in iTOL website (https://itol.embl.de/)^41^.

### Relative abundance profiling of MAGs

The abundance of MAGs constructed from 12 healthy adults was quantified by the *quant* function of *salmon* program (version 0.13.1) in metaWRAP pipeline^14^ and merged with taxonomy table. Phylum ‘Firmicutes_C’, ‘Firmicutes_A’ were converted to ‘Firmicutes’ to remove clade annotations. To avoid bias caused by absolute abundance different between samples, abundance of same phylum in the same host was added and their relative abundance was calculated and added based on the sampling point. Relative abundance was re-calculated to get relative abundances of phylum level on each sampling point. To compare the composition of top-5 abundant phylum between sampling points, minor phylum, which abundance was less than top-5, converted to ‘others’ in each sampling points. EARB containing community’s abundance table was further investigated at specie level and categorized into ‘EARB’, ‘top5’ (top-5 abundant species except EARB) and ‘others’. Lastly, to generate the relative abundance of *Enterobacteriaceae* and *Veillonellaceae*, family level taxonomy information of MAGs were categorized into ‘*Enterobacteriaceae*’, ‘*Veillonelellaceae*’ or ‘others’.

### Functional analysis of MAGs

Gene prediction using *Prodigal* (version 2.6.3)^42^ was conducted for further functional and community power analyses. Functions of predicted genes were annotated with KEGG using *hmmsearch* function of HMMER program (version 3.3.1)^43^ using pre-trained hidden Markov Models (prok90_kegg94). KEGG pathways/modules and the list of KEGG orthology (KO) terms was manually parsed from the KEGG database (release 103; 2022/07)^44^ and used for functional analysis. For comparison between non-EARB and EARB, samples that contain EARB were selected, and KO pathway and module table were generated by counting the number of KO genes for pathway or module in each MAG. A significant difference pathway or module was identified using a two-sided *Wilcoxon* rank-sum test with a confidence level of 0.95. Significant pathways were filtered based on the criteria - pathway gene coverage > 30% and fold change > 0.5 compared to the non-EARB. Significant modules were filtered based on the criteria - module gene coverage > 0.8 and fold change > 0.5 compared to the non-EARB.

### Community power analysis

To investigate the functional importance of EARB in a community, community power analysis was conducted. Genes of MAGs in the communities with EARB existing were predicted using prodigal (version 2.6.3) and KEGG annotated by HMMER (version 3.3.1). Therefore, KEGG enzyme count table was generated for each community. Functional uniqueness was measured by counting number of unique KEGG enzyme in a MAG when comparing others in the same community by leave-one-out approach. To calculate functional importance of a microbe, each KEGG enzyme counts of a microbe were divided by sum of the same enzyme counts of entire community member (proportion of KEGG enzyme for the microbe). The sum of proportion of all KEGG enzymes of a microbe was considered as community power score of the given microbe. Community power score of three categorized microbes (EARB, SARB, non-carrier) were also calculated.

### Co-enrichment analysis

Co-enrichment matrix was generated using mOTU relative abundance rarefied table downloaded from original paper^13^. First, we choose three mOTU species corresponding to EARB (Escherichia coli, Klebsiella oxytoca, Klebsiella pneumoniae). Four non-EARB species (*Bacteroides thetaiotaomicron*, *Bacteroides ovatus*, *Bilophila wadsworthia*, *Parabacteroides distasonis*) were manually picked based on high prevalence and high community power score for comparing. Co-enrichment correlation between each selected microbes’ abundance and other mOTUs’ were calculated by spearman test. mOTU species that annotated ambiguously (annotated as ‘motu_linkage_group’) and showed correlation between –0.3 < and < 0.3 with all selected species were excluded. The correlations that both microbes co-existed at least 5 communities and showed significant p-value (>0.05) were selected and plotted using a pheatmap R package. To count positively or negatively related mOTUs, filtering spearman correlation test by > 0.3 or < -0.3 for each comparing bacteria. To verify the relationship between relative abundance of EARB or non-EARB with mOTU richness, we sum up relative abundance of all EARB mOTUs or non-EARB in each sample. Then we counted mOTUs which is none zero in relative abundance in each sample. We calculated the sum of relative abundance as log2 and then divided the samples into three categories (Rare: log2 relative abundance sum < -10, Normal: -10 ≤ log2 relative abundance sum < -5, Rich: 5 ≤ log2 relative abundance sum). Correlation between relative abundance sum of EARB or non-EARB and number of none zero mOTUs (mOTU richness) was measured by spearman’s correlation coefficients.

### Tracking endogenous strains among metagenomic samples

To check genomic similarity between basic EARBs, we created an ANI matrix using fastANI^45^ with –matrix parameter with default options (version 1.33). For strain analysis, gene prediction was conducted for 12 EARB *E. coli* using *Prodigal*^42^. All predicted genes were concatenate then clustered based on similarity using CD-HIT (version 4.8.1)^46^ with -aS 0.9 -c 0.98 -n 10 - M 0 -d 0 -T 0 -G 0 parameter. From CD-HIT clustering information, extract the longest 20 homologous genes that present in all 12 *E. coli* strains with same length using cdbfasta (version 1.00). 20 homologous genes in most high-quality *E. coli* genome (ERR1995253 bin.3) were extracted and built as a reference by bwa (version 0.7.17-r1188) for variant call and SNP search. Variant call was done by following steps. Paired FASTQ files from EARB *E. coli* containing samples were mapped against previously built *E. coli* homologous gene reference. Generated bam file was sorted, index and mpileup using samtools (version 1.9)^47^. Variants were called using bcftools (version 1.9)^48^, and filtered to include only variants with a quality score of 20 or higher. Bam files was converted tabula form using sam2tsv version (d29b24f2b). Frequency of nucleotide in every variant point was counted using converted tsv file. Nucleotide bases which existed in variant position less than 5% were discarded because of the possibility of sequencing error. SNP ratio of each variant position was counted using basic R functions. For SNP analysis for AMR gene, we did the same process except using AMR gene from ERR1995248 bin.3 (host 5 & day 8) for reference because it is dominant *E. coli* of the host. Using dominant strain AMR gene for variant calling, we were able to find nucleotide ratio clustered into two in day 4 sample. Comparing variant position and SNP information between longitudinal data, we could track SNP ratio (SNP nucleotide frequency / total nucleotide frequency) of variant position through sampling points. To confirm the existence of two E. coli strain in host 5 and day 4 sample, all variant positions of 54 AMR gene reference host 5 at day 4 were investigated their sequence depth.

### Validation using different cohort dataset

To confirm the reproducibility of EARB in other antibiotic exposed human metagenome data, we downloaded raw data (i.e. FASQ files) of shotgun metagenomic samples, or isolate strain genome samples, from three different study dataset, such as RUTI-causing *E. coli* isolates (n=556), liver cirrhosis patient (n=592) and preterm infant (n=399). We constructed MAG from the raw data from liver cirrhosis patient and preterm infant data and search AMR gene using same pipeline, such as MetaWRAP pipeline, as we described above. For liver cirrhosis patients data, *Megahit*^39^ was used with -r parameter (for single-end input). For *E. coli* isolates data, we assembled contigs using *Megahit* then predicted genes using *prodigal* for AMR gene searching. We also investigated the *E. coli* isolates’ SNP profile of variant positions which are founded in SNP analysis between major strain and minor strain of host 5. We calculated AMR prevalence value and compared with healthy adults’ EARB. To verify the community power score of EARB in live cirrhosis and preterm infant data, we selected communities that contain EARB and more than 5 other MAG (liver cirrhosis = 28, preterm infant = 24 communities). Communities for healthy adults, liver cirrhosis patients and preterm infant were arranged by sampling point, MELD score (Model for End-Stage Liver Disease) and host day of life, respectively.

## Author contributions

JWB, NP, BS, NK, JN, AM, SS, J-I K, JWS, AK, and SL reviewed the manuscript and provided the critical comments. JWB and SL wrote the manuscript. JWB and SL performed the bioinformatics analysis. JWB and SL conceived the project.

## Funding

This work was supported by grants of the Basic Science Research Program (2021R1C1C1006336) and the Bio & Medical Technology Development Program (2021M3A9G8022959) of the Ministry of Science, ICT through the National Research Foundation; by a grant of the Korea Health Technology R&D Project through the Korea Health Industry Development Institute (KHIDI), funded by the Ministry of Health & Welfare (HR22C141105), South Korea; by the “Korea National Institute of Health (KNIH)” research project (2024-ER2108-00; 2024-ER0608-00); and also by a GIST Research Institute (GRI) GIST-MIT research Collaboration grant by the GIST in 2024.

## Extended Figure Legends

**Extended Fig. 1.**
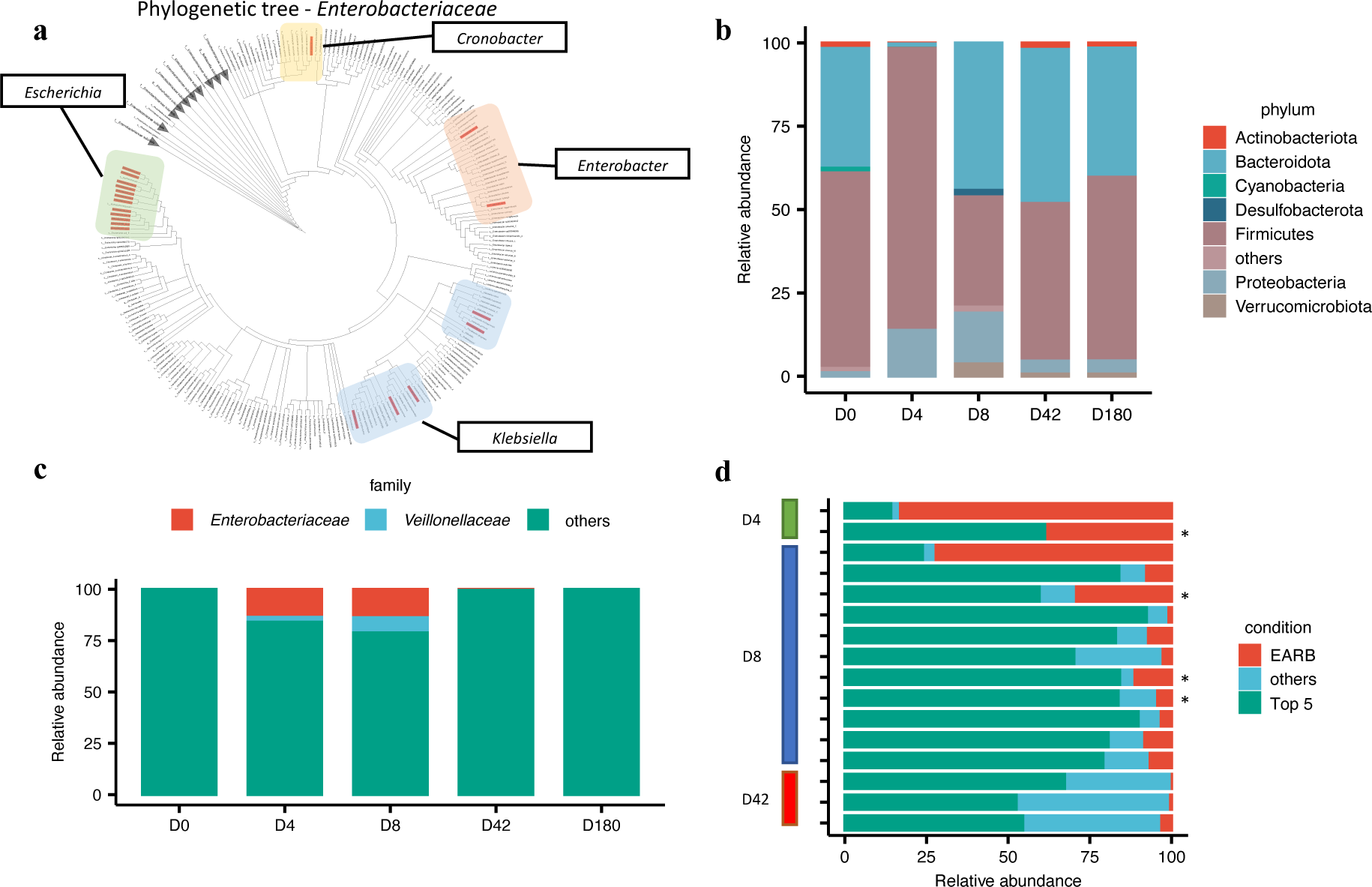
Taxonomy and functional comparative analysis of MAGs revealed unique characteristics of EARB versus other bacteria in metagenomic samples. **a,** Phylogenetic tree of MAGs and *Enterobacteriaceae* reference genomes. All EARB, highlighted in red within the tree, are distributed across four genera within this family. **b,** The relative abundance of the top five phyla and other taxa of given MAGs at each time point. **c,** Co-enrichment of *Enterobacteriaceae* and *Veillonellaceae* shown as the relative abundance profiles. The abundance of a given family was calculated as the sum of the quantified abundance of given MAG bins using metaWRAP. **d,** The relative abundance profiles of EARB strains and other MAGs (only samples where EARB strains existed [n = 16] are shown). Asterisk marks (*) indicate samples with two EARB strains identified.

## Supplementary Figure Legends

**Supplementary Fig. 1.**
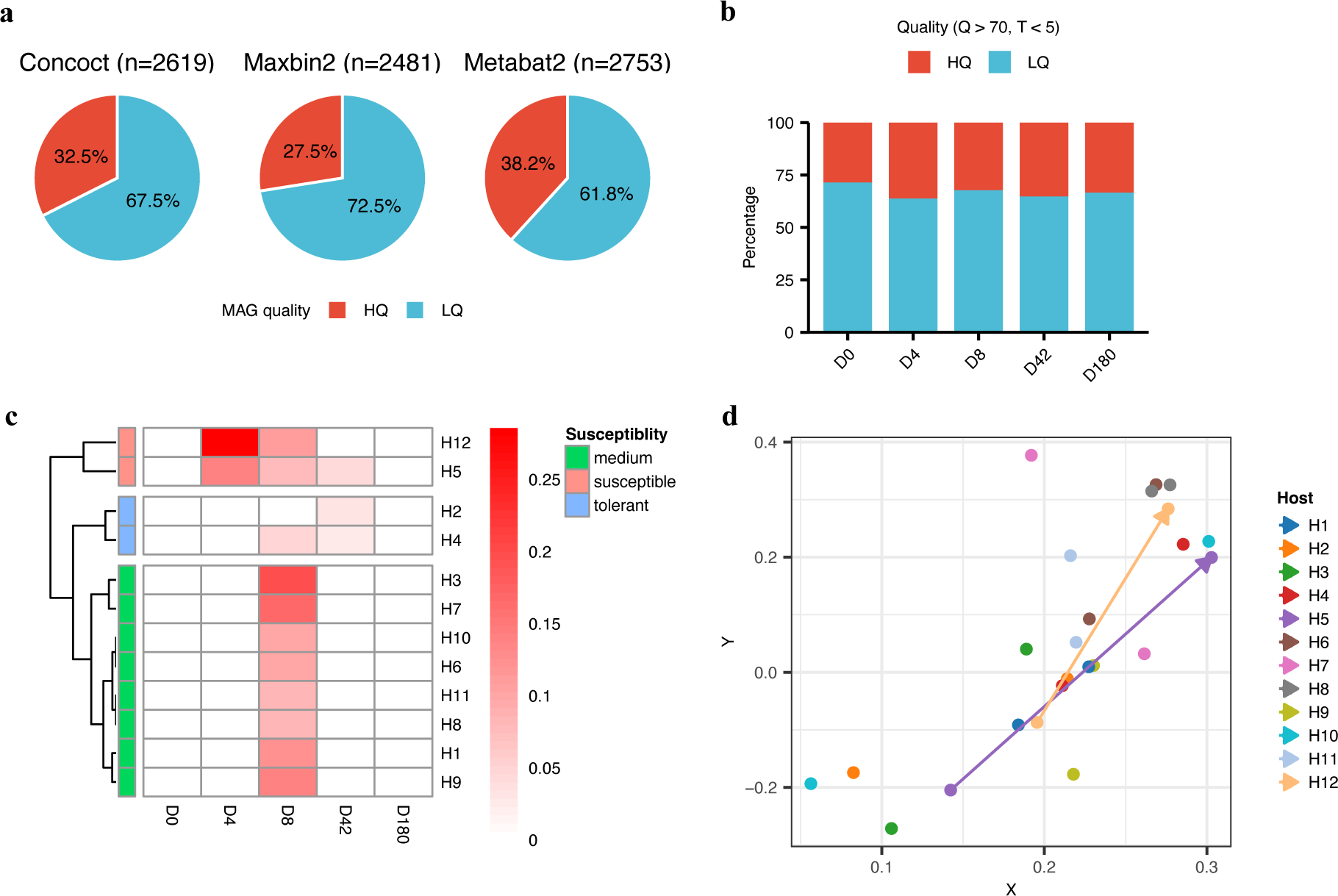
Statistics of high-quality MAGs from three different binning tools and individual-specific changes in the resistome detected among high-quality metagenome-assembled genomes (MAGs) across different time points before and after antibiotic treatments. **A**, The percentage of high-quality (HQ) and low-quality (LQ) MAGs identified in the three different binning tools, Concoct, Maxbin2, and Metabat2. High-quality thresholds were based on 1) completeness (Q) greater than 70% and 2) contamination (T) less than 5%. **b**, The percentage of high-quality (HQ) and low-quality (LQ) MAGs for each time point over five sampling points (day [D] 0, 4, 8, 42, and 180). **c**, The fraction of EARB MAGs in the total MAGs in each metagenome sample. For brevity, we imputed fractions of three missing samples as zeros (host [H] 1, 8, 10 at day-4). **d**, Bray-Curtis distances on principal component analysis plots based on the relative abundance of all detected microbes by metagenomic-based operational taxonomic units, at D0 and D180. Arrows indicate samples of susceptible hosts (H5, H12), connecting them from D0 to D180. Samples from the above-mentioned hosts showed the longest distances between D0 and D180 among other hosts, except H10.

**Supplementary Fig. 2.**
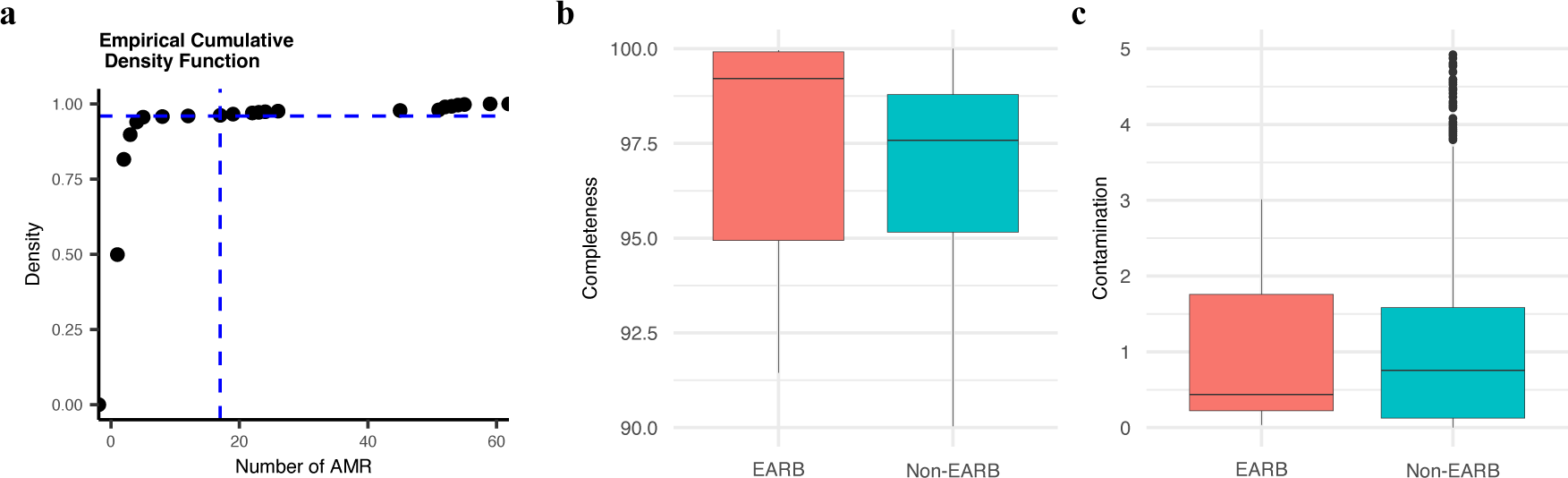
the determination of high-quality EARB MAGs based on the cumulative distribution of the number of antibiotic-resistance (AMR) genes of MAGs and their binning quality scores. **a**, The empirical cumulative density function (eCDF) of all MAGs with an AMR gene (n = 499). The horizontal blue dotted line corresponds to 0.96 and the vertical line corresponds to 17. **b–c**, binning quality score calculated using CheckM for EARB and non-EARB strains.

**Supplementary Fig. 3.**
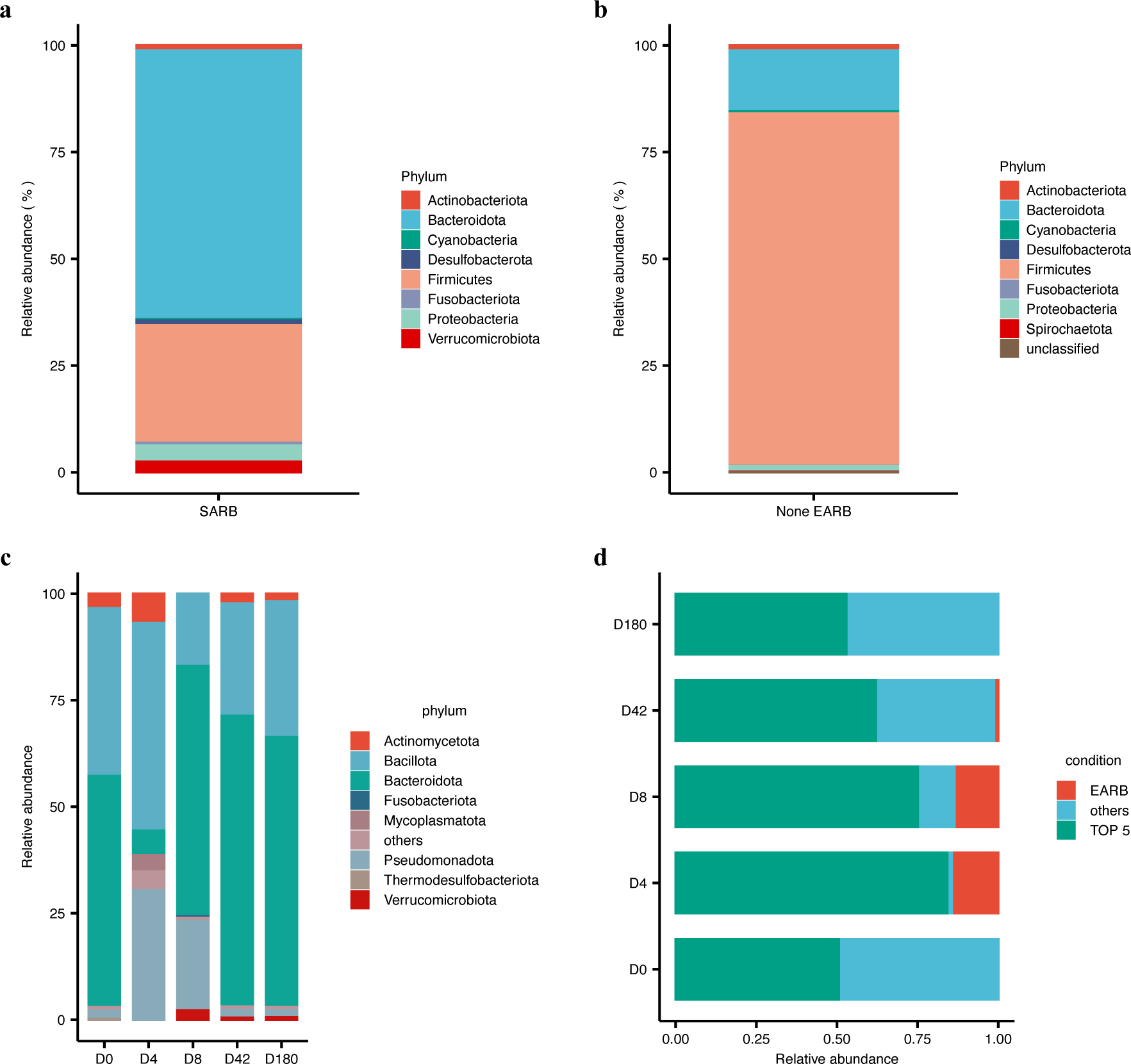
The average relative abundances of (**a**) SARB strains and (**b**) non- EARB strains. Average relative abundances of SARB (n = 479) and non-EARB (n = 859) strains are shown at the phylum level. **c**, Relative abundance of the core microbiome (i.e., top 5 highly abundant phylum) in each condition. The abundance of each phylum was calculated using Kraken2. Other non-core microbiome components are shown as “others”. **d**, Summary of relative abundance of EARB and core microbiome in all samples.

**Supplementary Fig. 4.**
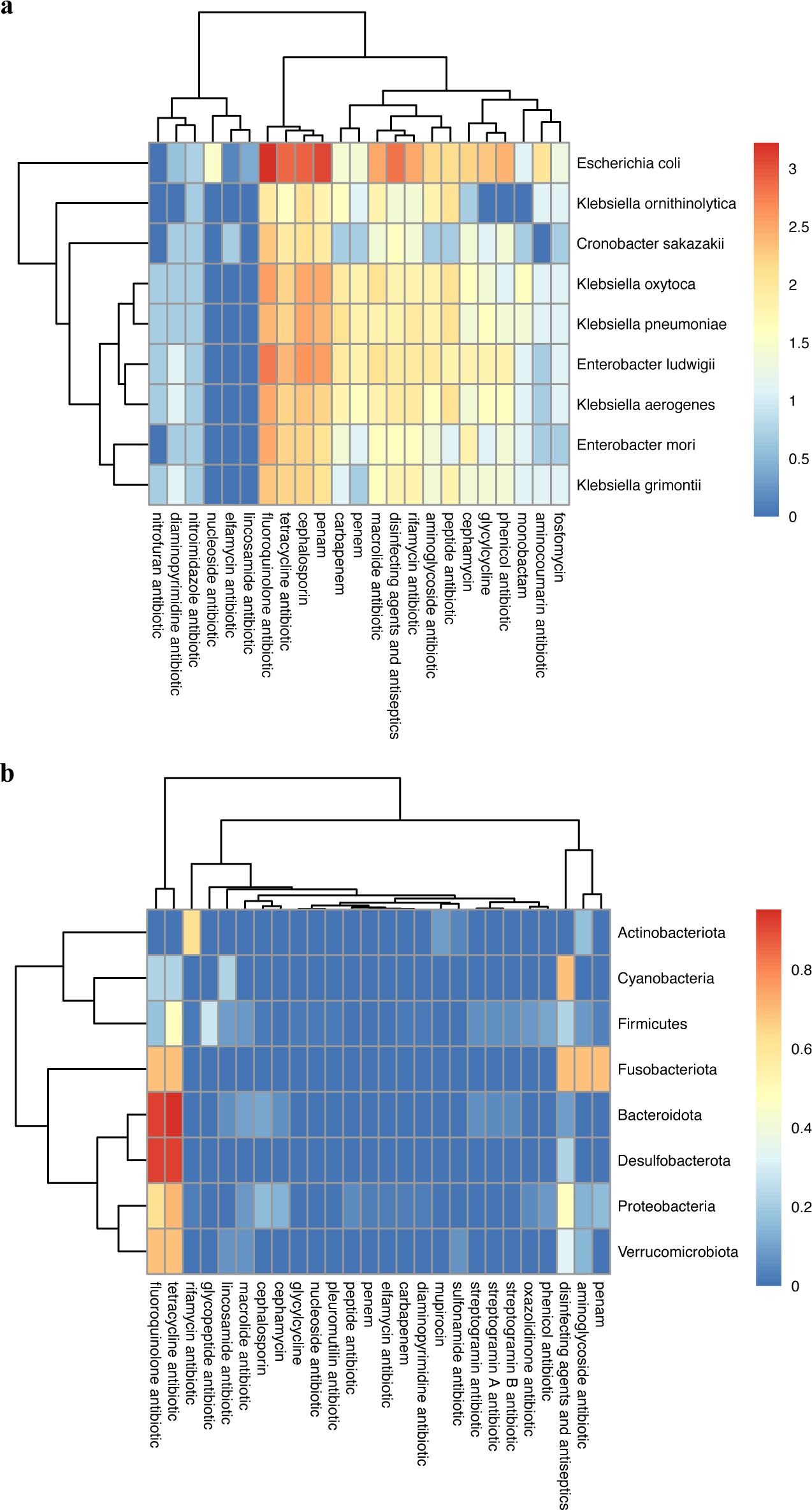
Frequency of resistant drug class of given taxon of AMR bacteria. **a–b**, The proportion of the AMR genes of a given resistant drug class per specific taxonomic unit is shown for (**a**) EARB (plotted at the species level) and (**b**) SARB (plotted at the phylum level).

**Supplementary Fig. 5.**
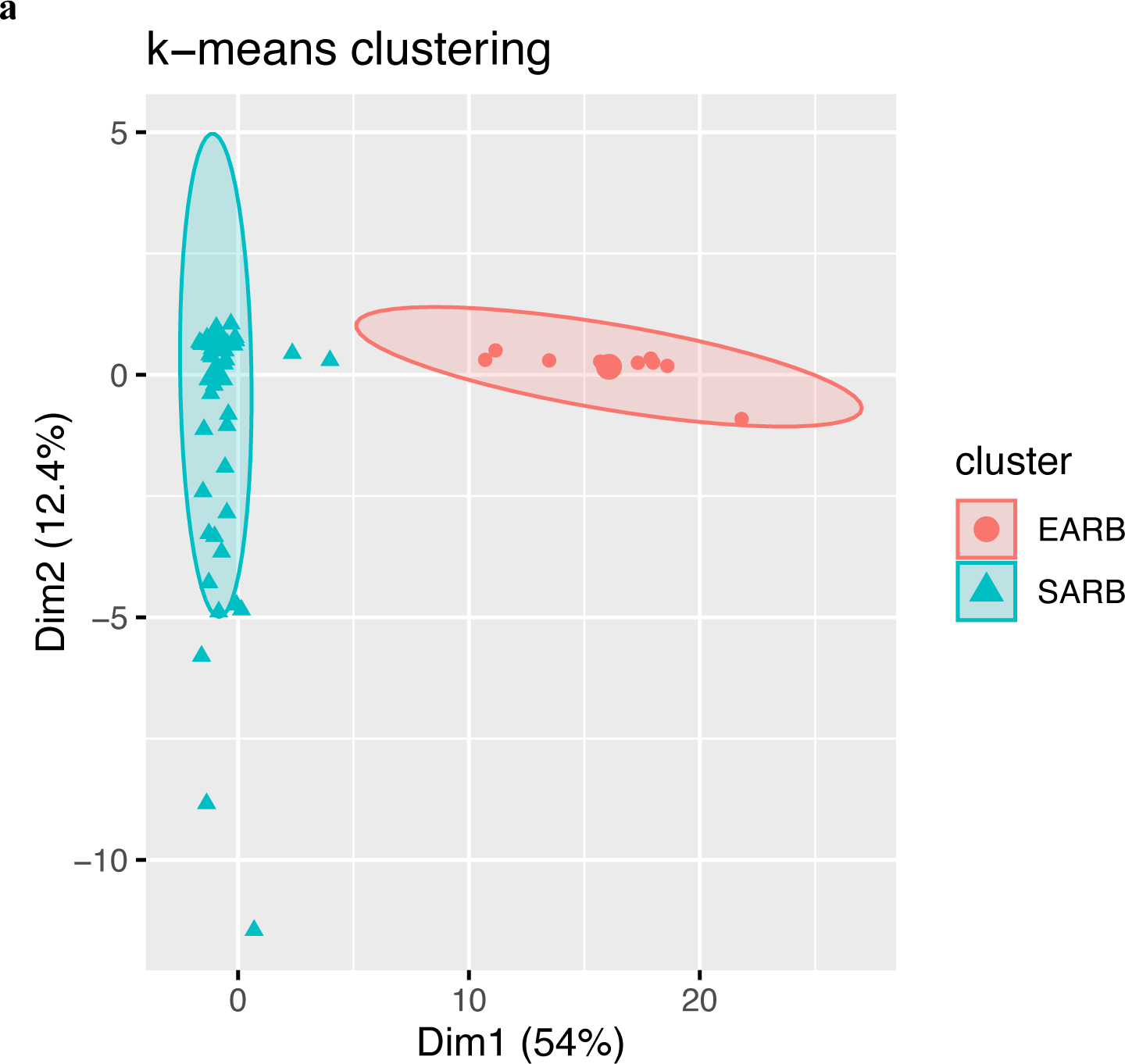
k-means clustering using resistome of EARB and SARB. Relative frequency of drug classes, frequency of AMR genes which confer resistance against to specific drug class / sum of all drug classes appearance in AMR genes, was calculated by Resistance Gene Identifier (RGI) result in species level.

**Supplementary Fig. 6.**
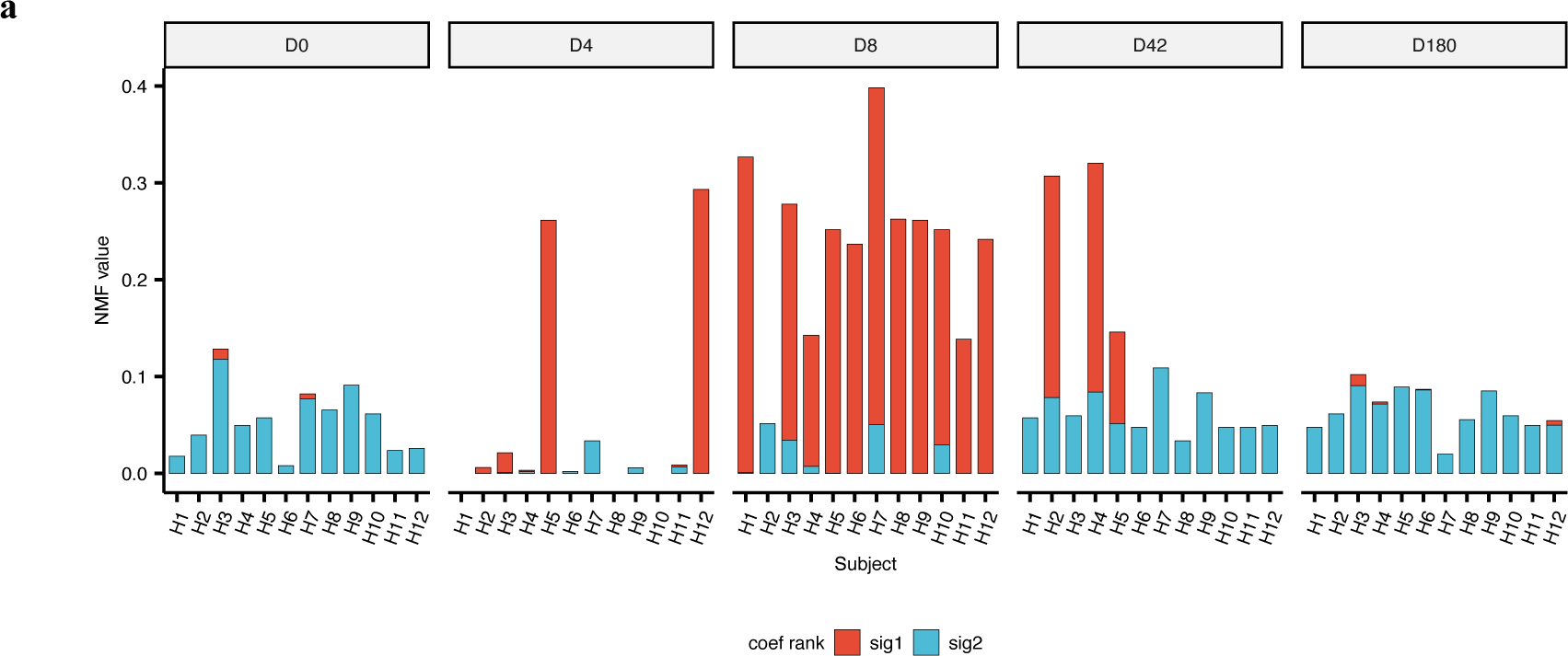
Non-negative matrix factorization (NMF) weight values of two signatures over five different time points by different hosts. We observed that the EARB-like signature (sig1) was highly enriched after antibiotic treatments (e.g. day [D]4 and D8), whereas the SARB-like signature (sig2) was evenly distributed in different time points, including D0.

**Supplementary Fig. 7.**
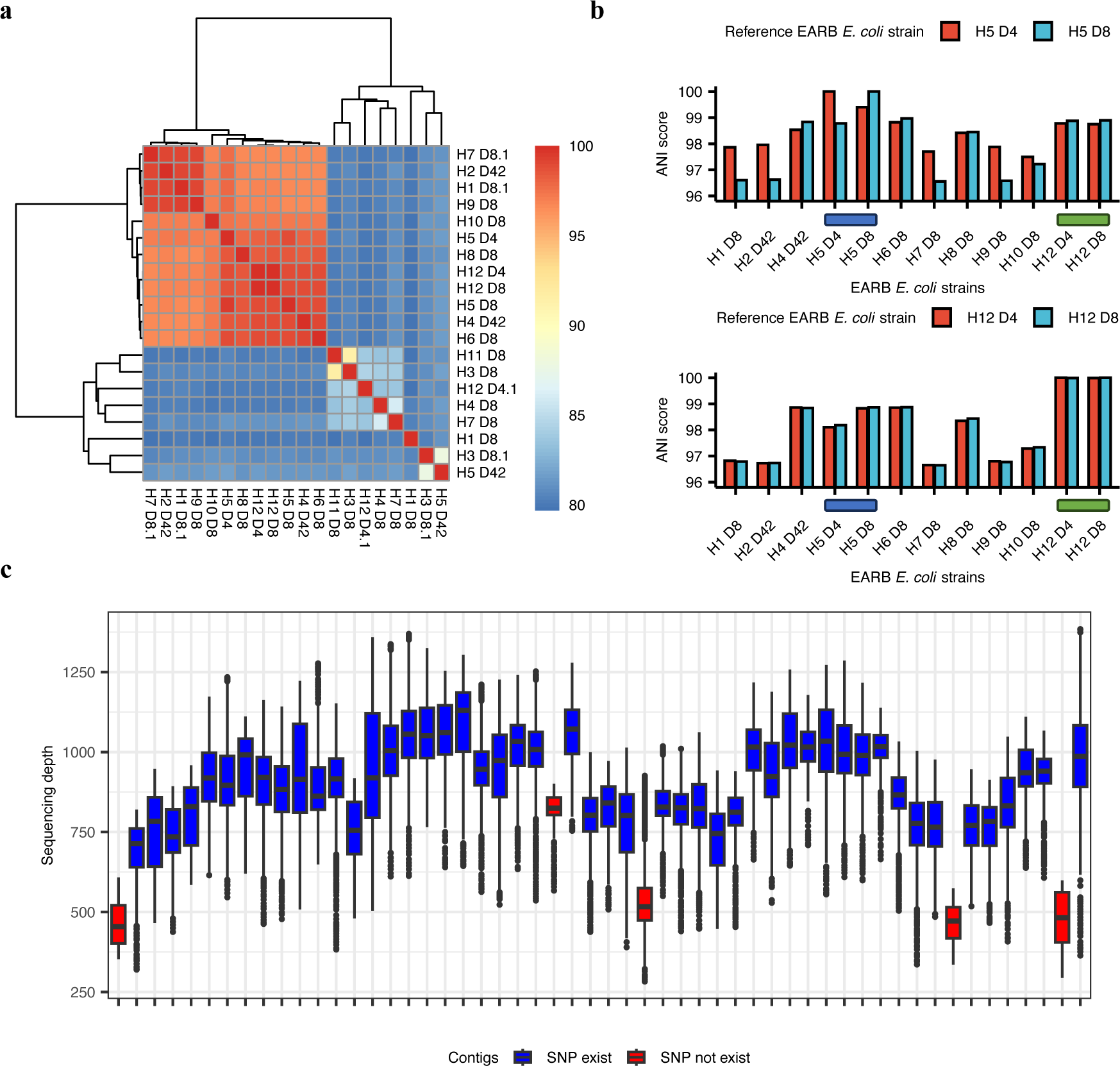
Diversity of EARB strains identified from ANI scores and SNP profiles. **a**, Average nucleotide identity (ANI) score matrix of EARB MAGs (n = 20). **b**, Comparison of EARB *E. coli* strain ANI scores with two specific strains identified from the same host (H5, H12). Blue and Green bar indicate two strains from the same host. **c**, Sequence depths of host (H)5, day (D)4 samples for 54 AMR genes found in the H5 D8 sample, which is the sample in which a major strain of EARB *E. coli* species was dominant. When we performed variant calls using D4 samples based on AMR genes identified in D8 samples, we found some genes with SNPs detected (blue) showed high sequencing depths, whereas genes having no SNPs showed relatively low sequencing depths in a given D4 sample.

**Supplementary Fig. 8.**
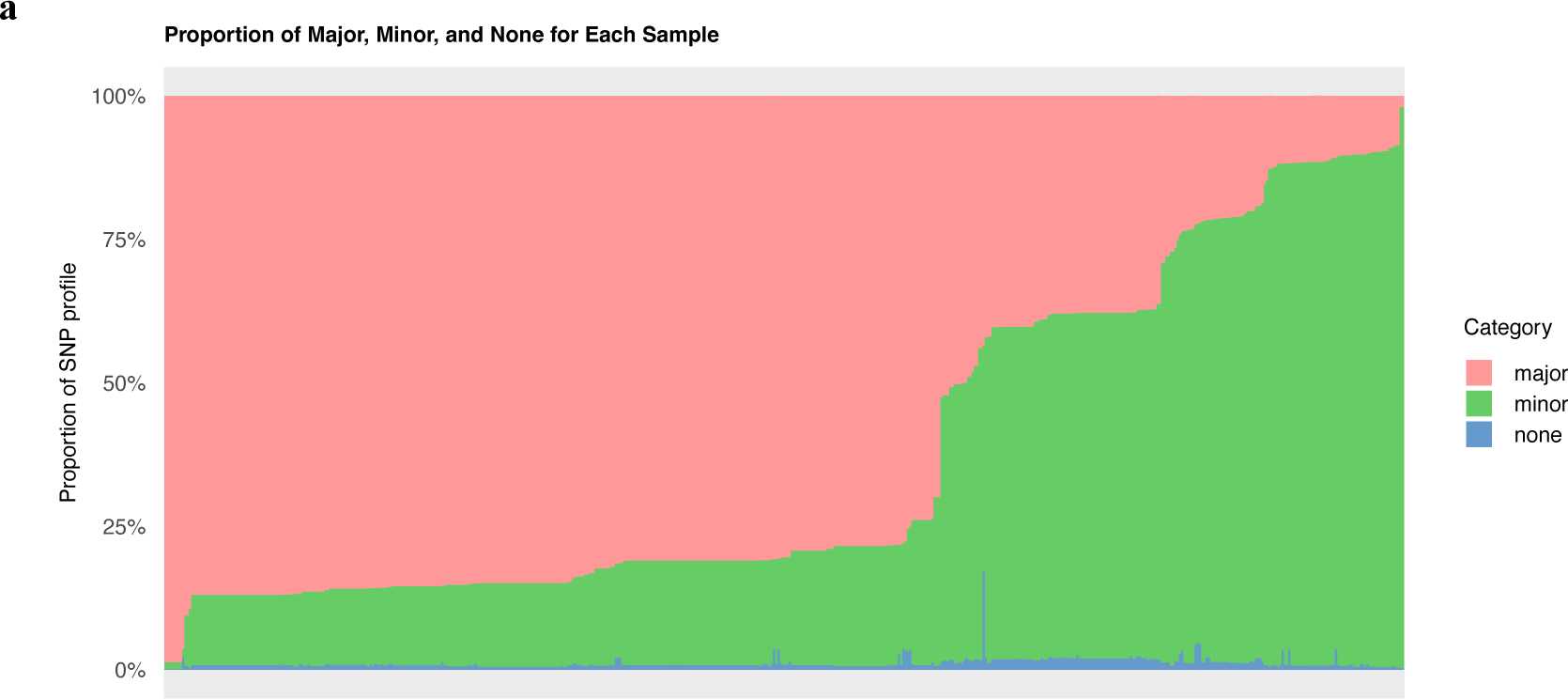
Proportion of major, minor, and none for all variant positions (n=1841) of RUTI *E. coli* isolates. If the most frequent nucleotide of the variant position is equal to the host 5 major *E. coli* strain’s allele, we categorized that variant position as major. About 65% of isolates (n=360) contain major allele more than half of the variant positions.

## Supplementary Table Legends

Table S1. Binning quality and taxonomic information of 1,358 metagenome-assembled genomes (MAGs) obtained from three binning tools (**Supplementary Data 1**)

Table S2. Annotation of antimicrobial resistance (AMR) genes of high-quality MAGs assembled from 12 healthy adults treated with antibiotic cocktails

Table S3. Spearman’s correlations between the abundances of microbes with the highest community power scores and other microbes detected using metagenomic-based operational taxonomic unit (mOTU) markers. Microbes that showed correlations between –0.3 and 0.3 with all selected species were excluded. The correlations in which both microbes co-existed in at least five communities and showed significant *p*-values (> 0.05) were selected.

Table S4. Merged core gene SNP analysis result from day 0 to day 8

Table S5. Merged AMR gene SNP analysis result from day 0 to day 8

Table S6. Annotation of antimicrobial resistance (AMR) genes of RUTI-causing *E. coli* isolates

Table S7. Binning quality and taxonomic information of 2,092 MAGs assembled from shotgun metagenome data of liver cirrhosis patients (**Supplementary Data 2**)

Table S8. Binning quality and taxonomic information of 1,150 MAGs assembled from shotgun metagenome data of preterm infants (**Supplementary Data 3**)

## Supplementary Data Legends

**Supplementary Data 1.** Sequence file of metagenome-assembled genomes (MAGs) assembled from shotgun metagenomic data from 12 healthy adults treated with antibiotic cocktails

**Supplementary Data 2.** Sequence file of metagenome-assembled genomes (MAGs) assembled from shotgun metagenome data from patients with liver cirrhosis (PRJEB38481).

**Supplementary Data 3**. Sequence file of metagenome-assembled genomes (MAGs) assembled from shotgun metagenome data from preterm infants (PRJNA301903)

## Notes

### Competing Interest Statement

The authors have declared no competing interest.

