## Supplementary figures and images for "Extensively acquired antimicrobial resistant bacteria restructure the individual microbial community in post-antibiotic conditions"

### Extended Figure 1

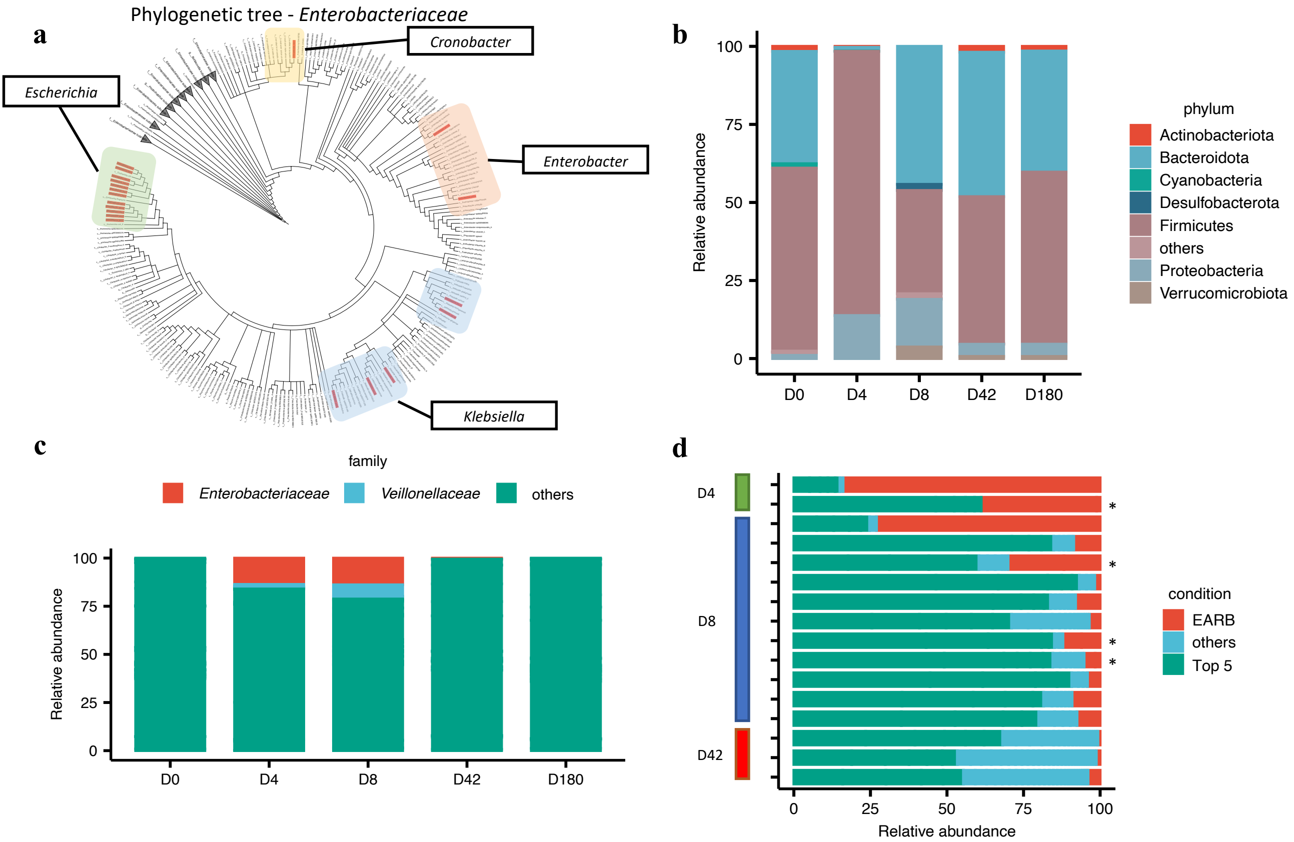

### Supplementary Figure 1

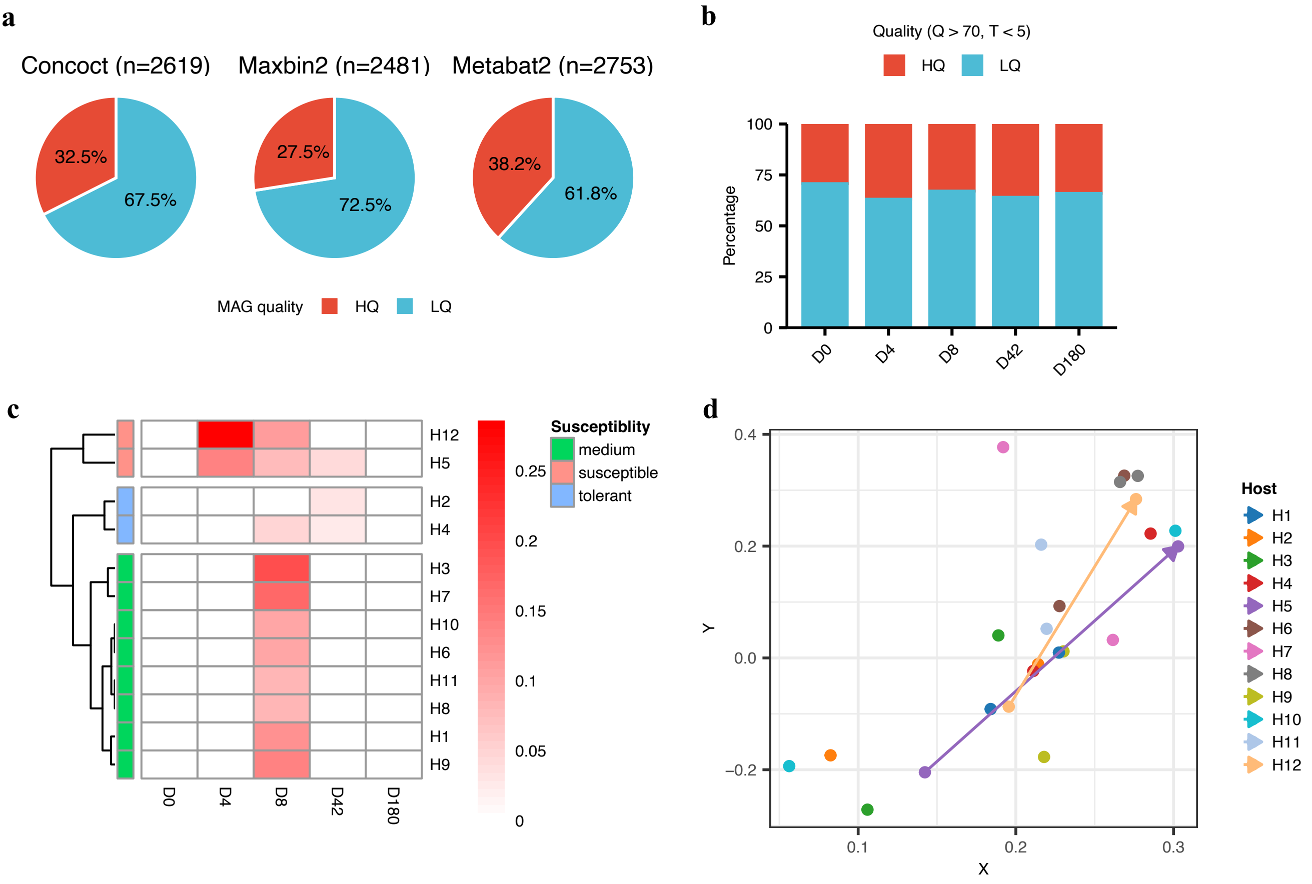

### Supplementary Figure 2

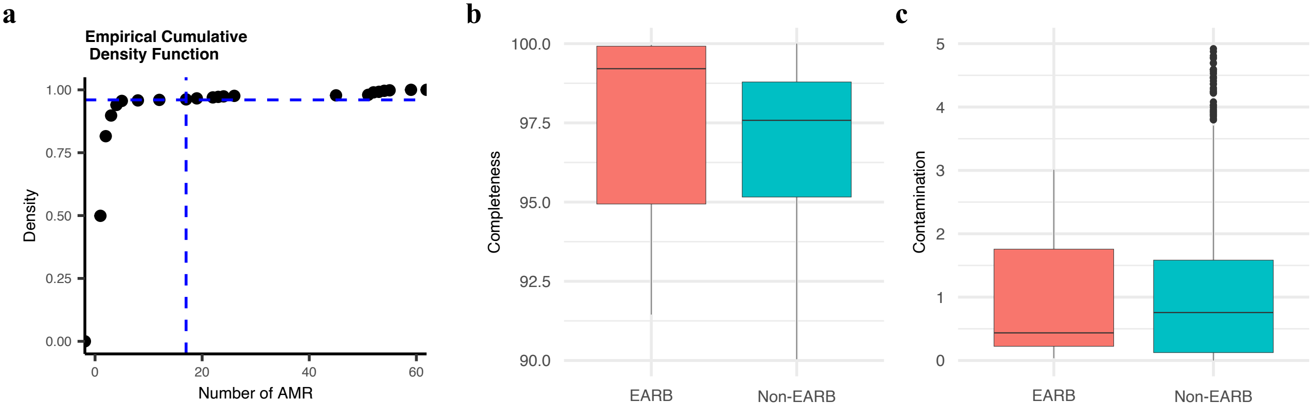

### Supplementary Figure 3

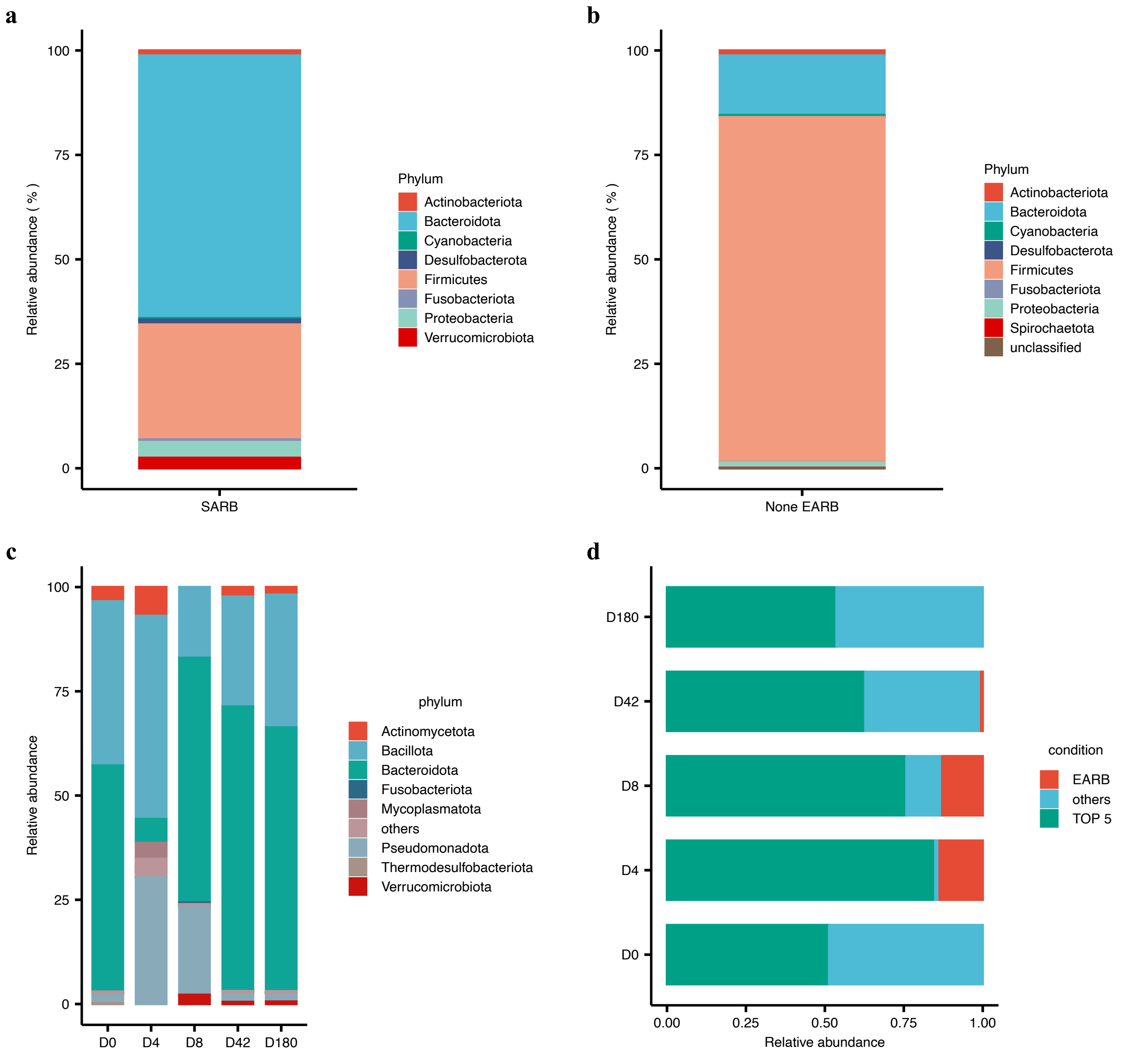

### Supplementary Figure 4

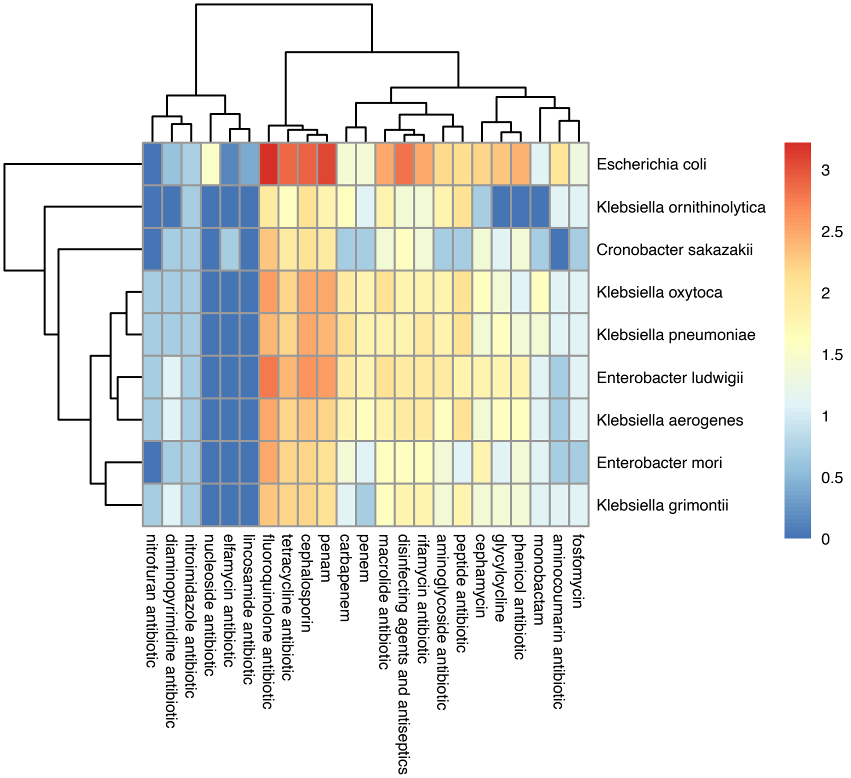

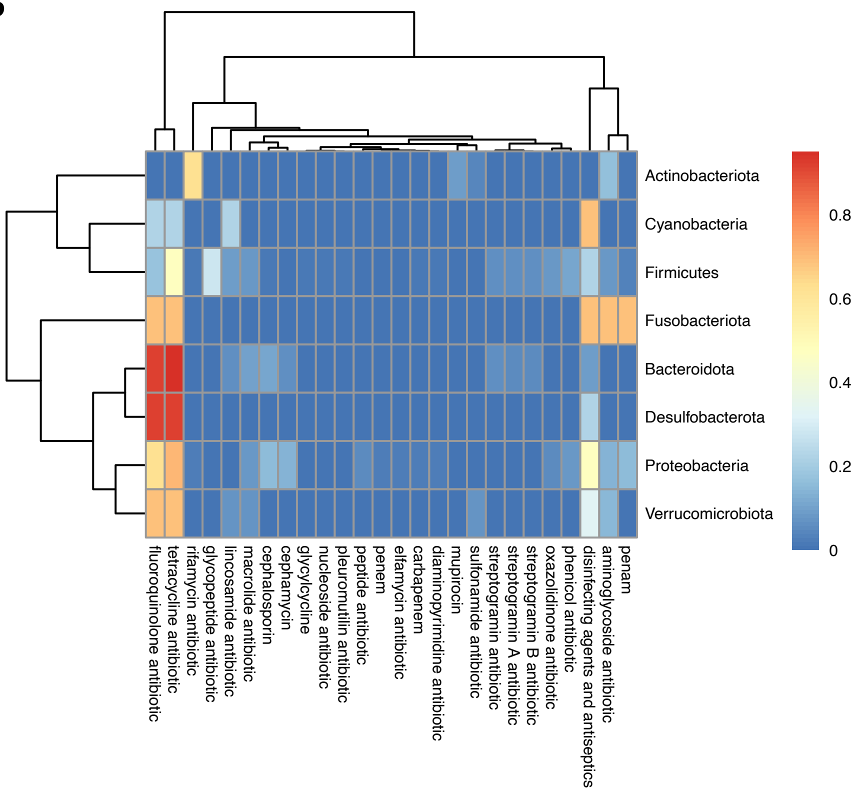

### Supplementary Figure 5

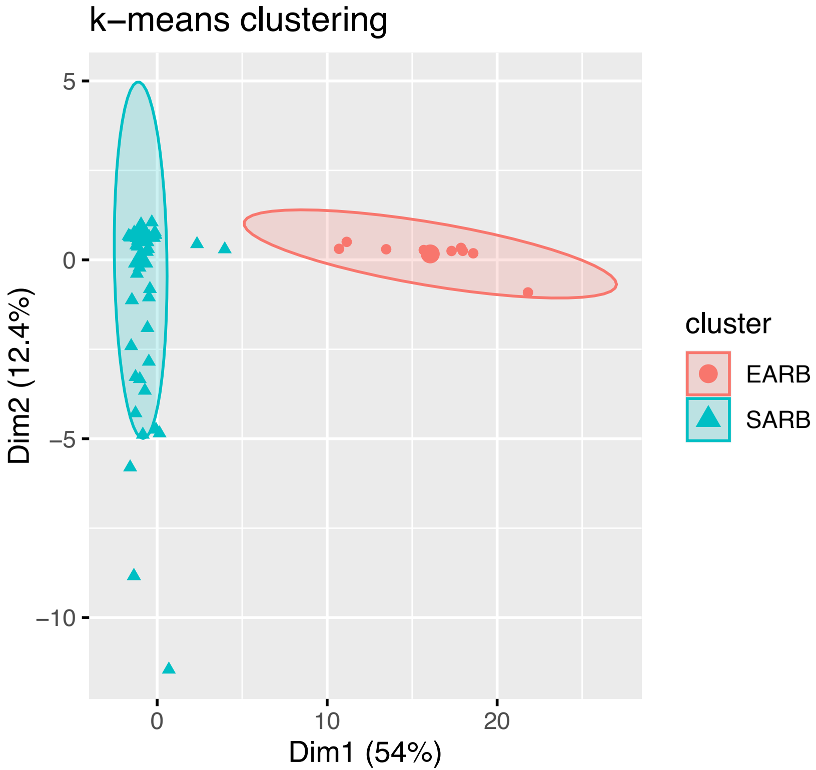

### Supplementary Figure 6

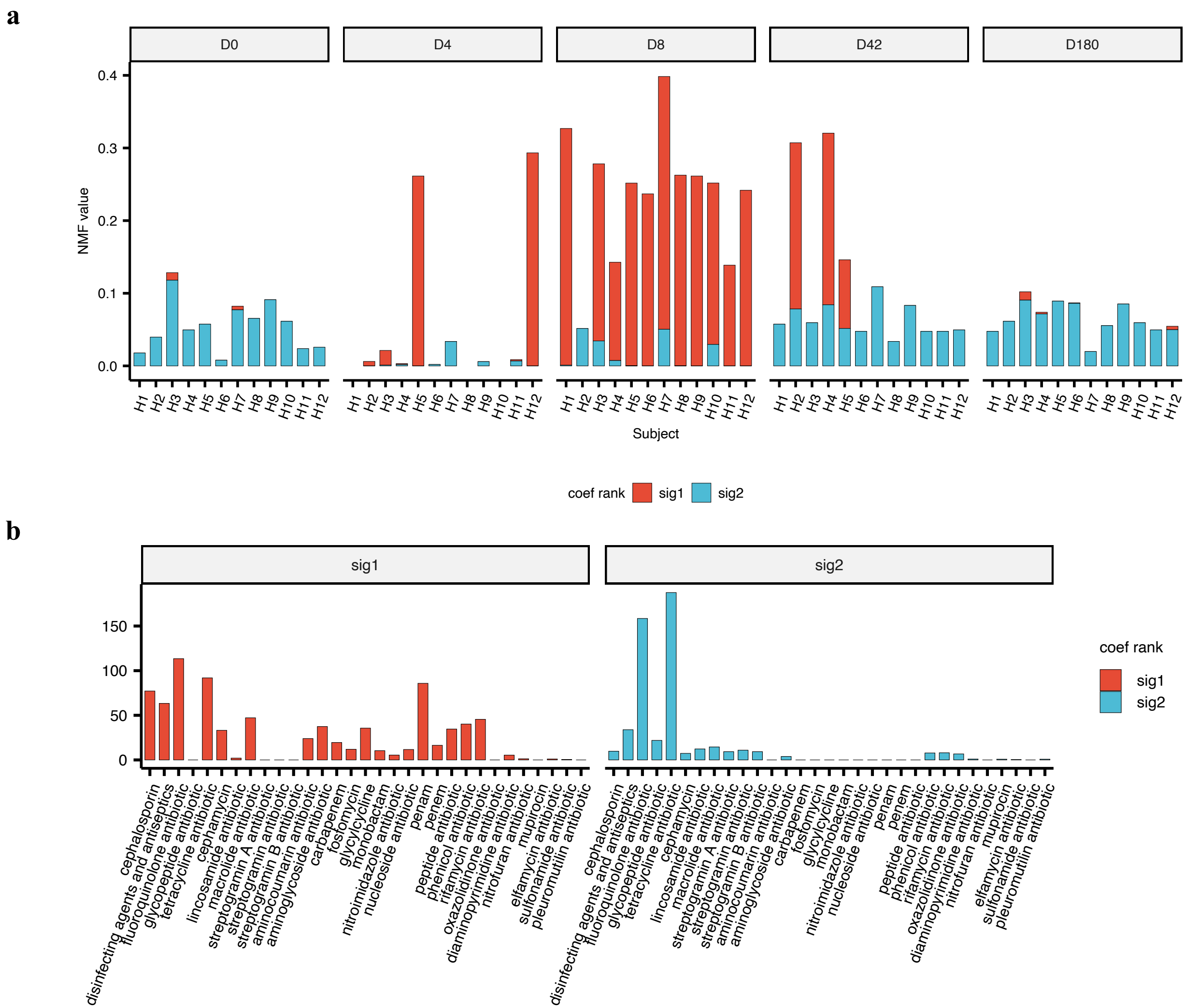

### Supplementary Figure 7

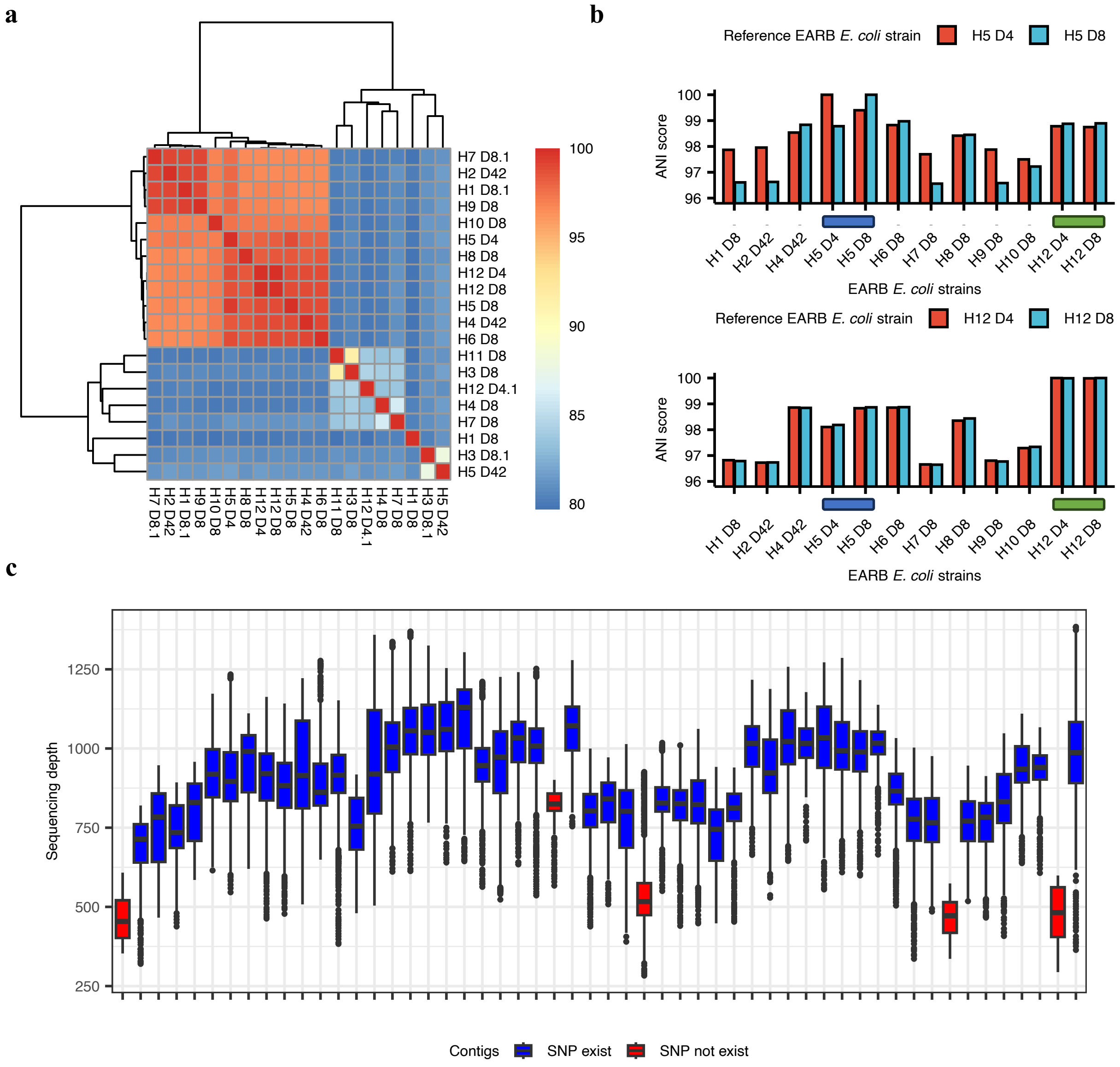

### Supplementary Figure 8

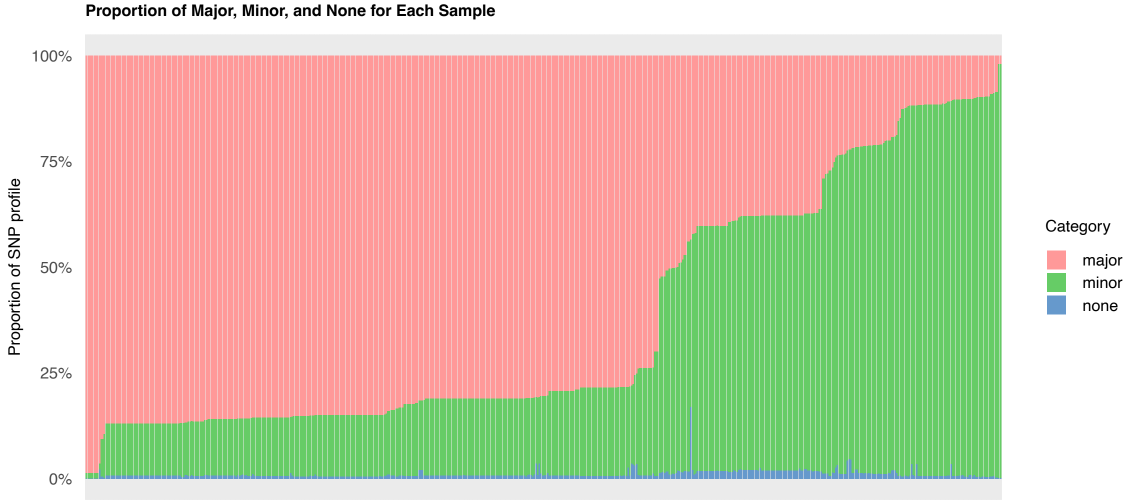
